# A single-component optogenetic toolkit for programmable control of microtubule

**DOI:** 10.1101/2025.10.31.685931

**Authors:** Guolin Ma, Xiaoxuan Liu, Tien-Hung Lan, Tam Duong, Megan Chiu, Danica Du, David J. Reiner, Yun Huang, Yubin Zhou

## Abstract

Microtubules (MTs) form dynamic cytoskeletal scaffolds essential for intracellular transport, organelle positioning, and spatial organization of signaling. Their architecture and function are continuously remodeled through the concerted actions of microtubule-associated proteins (MAPs), post-translational modifications (PTMs), and molecular motors. To precisely interrogate these processes in living systems, we developed a genetically encoded optogenetic toolkit for spatiotemporal control of MT organization and dynamics. By replacing native multimerization motifs with a blue light-responsive oligoermization domain, we have engineered single-component probes, OptoMT and OptoTIP, that reversibly label MT polymers or track plus-ends with tunable kinetics from seconds to minutes. When coupled to enzymatic effectors, these modules enable localized tubulin acetylation or detyrosination, directly linking PTMs to MT stability. We further engineered OptoMotor, a light-activatable kinesin platform that reconstitutes tail-dependent cargo transport along MTs, and OptoSAW, a light-triggered severing actuator for controlled MT disassembly. Using these tools, we reveal how local MT integrity governs lysosomal trafficking and ER-associated signaling dynamics. Collectively, this versatile single-component toolkit bridges molecular design with cytoskeletal function, offering new avenues to illuminate how dynamic cytoskeletal architectures coordinate intracellular organization, transport, and signaling.

## INTRODUCTION

Microtubules (MTs) form a dynamic and polarized cytoskeletal network that support a broad spectrum of cellular processes, from directed intracellular transport and cell division to signal transduction and organelle positioning [1, 2]. These varied roles dependent on a network microtubule-associated proteins (MAPs) that encompass molecular motors, adaptor proteins, severing enzymes, and regulators of post-translational modifications (PTMs) [3, 4]. While decades of biochemical and structural studies have delineated many of core mechanisms underlying MT assembly and turnover [5, 6], they often fall short to capture the highly localized and transient regulation that occurs in living cells, where MAPs tend to operate in concert and context-dependently.

Optogenetic systems have transformed cell biology by enabling precise, reversible, and non-invasive control of protein activity, localization, and interaction dynamics in real time [7–14]. Despite their success across signaling, transcriptional, and cytoskeletal pathways, harnessing optogenetic control of the MT network remains comparatively nascent, in part owing to the structural complexity of MTs and the functional diversity of MAPs that mediate their regulation [15–18]. Bridging this gap requires a versatile optogenetic platform that can interrogate MT dynamics across multiple regulatory layers, ranging from structural organization and PTM modulation to cargo transport and filament remodeling, under physiologically relevant conditions.

A fundamental prerequisite for such an approach is the ability to visualize and modulate MT architecture with minimal perturbation. Conventional methods such as tubulin overexpression, immunostaining, or chemical labeling often perturb polymerization kinetics or are incompatible with live-cell imaging [19–21]. In contrast, a genetically encoded photoswtichable system that transiently associates with MT lattices or plus-ends and can be coupled to effector domains offers the promise of interrogating MT behaviour and MAP function with sub-cellular resolution. Such a platform could, in principle, be adapted to manipulate enzymatic modifications of tubulin or to locally recruit effector proteins in response to light, thus extending optical control to the biochemical regulation of MT stability and organization.

Beyond structural support, MTs serve as the tracks for intracellular cargo movementtransport, a process driven predominantly by the kinesin family of MT-based motors [22]. For example, kinesin-1 is a prototypical plus-end-directed motor whose processivity depends on head-dimerization via its coiled-coil stalk and cargo engagement of cargo via through its tail domain [22, 23]. While optogenetic approaches have been developed to tether motors to cargo through light-induced heterodimerization using two-component systems [24, 25], they typically bypass native regulatory mechanisms such as motor dimerization and tail-dependent cargo selection. Harnessing optogenetic control that preserves, rather than circumvents, native kinesin gating offers a powerful route to dissect how motor assembly and regulation govern directional transport in live cells.

Third, equally important dimension to MT biology is filament remodelling, for example by severing enzymes such as spastin and katanin, which catalyse MT fragmentation to support network plasticity. These AAA+ ATPases assemble into hexameric rings that translocate along MTs and extract tubulin subunits through ATP hydrolysis, thereby generating new filament ends that promote network reorganisation and turnover [26–28]. Although in vitro and structural studies have provided mechanistic insight [29], the ability to control severing events with spatiotemporal precision in living cells remains limited. Achieving such control would enable direct interrogation of how MT remodelling influences organelle trafficking, calcium signalling and inter-organelle communication.

Here, we present a suite of optogenetic tools that enable reversible and modular control over distinct facets of the MT cytoskeleton. These include light-inducible probes for labeling and tracking MT filaments (OptoMT) and plus-end dynamics (OptoTIP), systems for localized modulation of tubulin post-translational modifications, optically controlled kinesin motors (OptoMotor) for tunable cargo transport, and photoactivatable severing modules (OptoSAW) for manipulating filament turnover in living cells. Together, this integrated toolkit establishes a unified platform for dissecting the structural, transport, and remodeling mechanisms that underpin MT organization, thereby bridging molecular interactions to cellular-scale dynamics and providing new avenues for exploring cytoskeletal function in physiology and disease.

## RESULTS

### Design of optogenetic probes for reversible labeling of MT cytoskeleton

To achieve reversible MT labeling with minimal perturbation to its dynamics, we exploited light-induced oligomerization of the N-terminal photolyase homologous region (PHR; residues 1-498) of *Arabidopsis* cryptochrome 2 (CRY2) [30–33] to cluster weak tubulin-binding motifs, thereby shifting their binding equilibrium toward stronger MT association in a light-dependent manner (**Fig. 1a**). We generated a panel of chimeric constructs by fusing CRY2 to tubulin-binding domains derived from four well-characterized MT binders, including kinesin (KIF5A), cytoplasmic linker region of 170 kDa (CLIP170), end-binding protein 1 (EB1), and calmodulin regulated spectrin associated protein 1 (CAMSAP1) (**Fig. 1b**) [1, 34–40]. In the dark, the designed hybrid protein is anticipated to be evenly distributed in the cytosol, whereas blue light stimulation at 470 nm could trigger rapid and reversible tracking of MT network (**Fig. 1a**). Among all tested constructs, CRY2-CLIP170_129-350_ (designagted “OptoMT”) yielded the least basal activity while exhibiting the robust MT labeling, characterized by a high signal-to-noise ratio and excellent reversibility (**Fig. 1c** and **Supplementary Videos 1-2**). Localized blue light stimulation of a defined region of interest (ROI) induced confined MT labeling within and immediately adjacent to the illuminated area, demonstrating subcellular spatial precision (**Fig. 1d** and **Supplementary Video 3)**. OptoMT exhibited rapid kinetics, with an activation half-life (*t*_1/2, ON_) of 10 s and a decay half-life (*t*_1/2, OFF_) of 210 s following cessation of illumination, making it suitable for dynamic and reversible live-cell imaging (**Fig. 1e**, **Supplementary Fig. 1**, and **Supplementary Table 1**). Labeling specificity was further validated by immunostaining in HeLa cells expressing mCherry (mCh)-OptoMT with an anti-α-tubulin antibody. Fixed-cell analysis revealed tight colocalization of OptoMT with the endogenous MT network (**Fig. 1f**).

**Figure 1.**
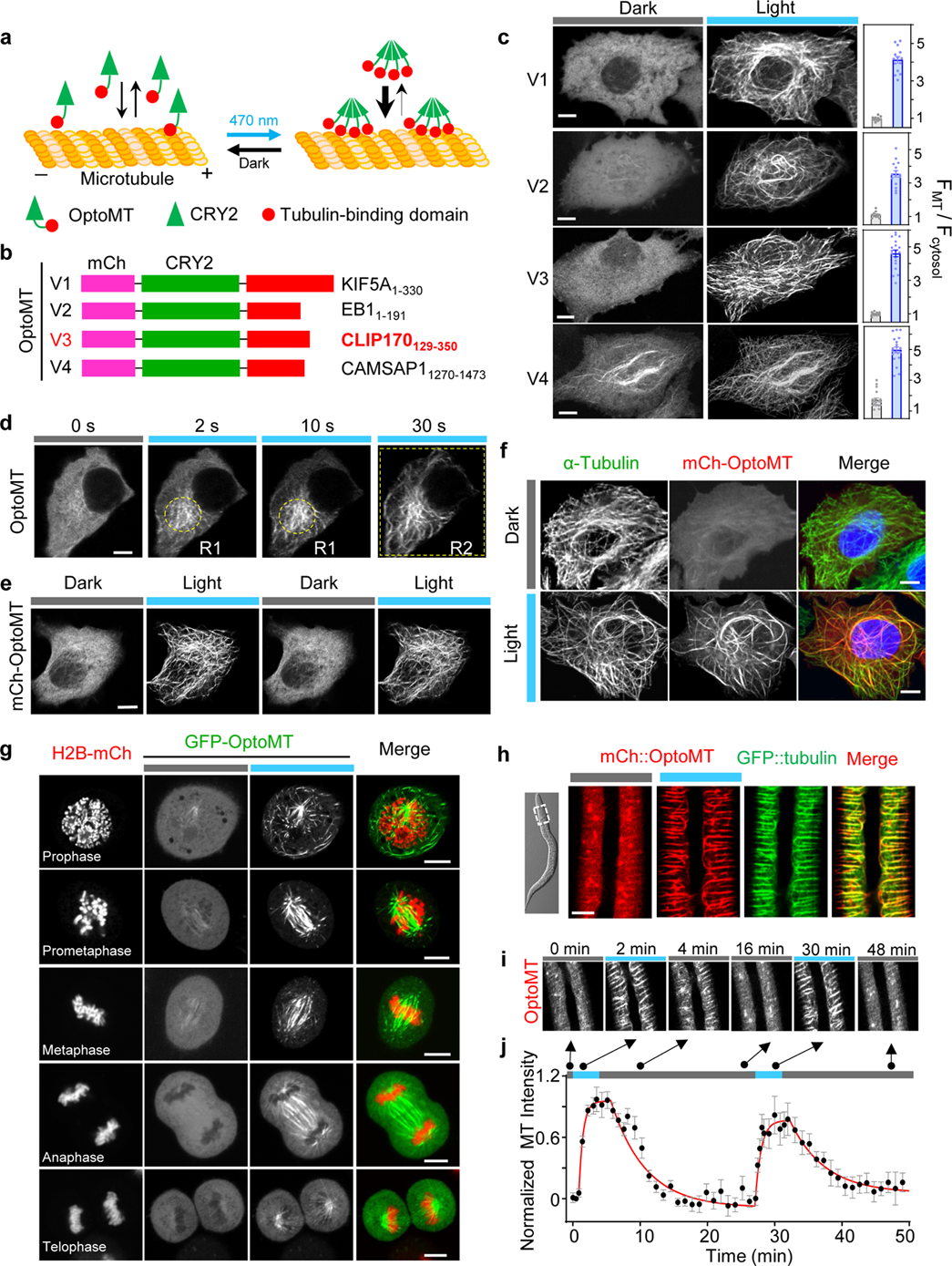
OptoMT for photo-inducible visualization of microtubule and mitosis in living cells. Scale bar, 5 μm. Error bars denote sem. **(a)** Schematic of OptoMT design and its light-induced association with MT. **(b)** Domain architecture of OptoMT variants. The photosensitive module CRY2 (aa 1-498) was fused to tubulin-binding domains derived from Kinesin, EB1, CLIP170, or CAMSAP1. The optimal construct (variant 3, highlighted in red) exhibited minimal basal activity in the dark but strong MT binding upon blue light stimulation. **(c)** Confocal images of HeLa cells expressing the indicated mCh-OptoMT variants with or without blue light exposure (indicated by blue bars). Right, quantification of normalized MT-to-cytosol fluorescence intensity ratio before and after 1 s light stimulation (470 nm, 40 μW/mm²). n = 24 cells from three independent biological replicates. Also see **Supplementary Videos 1-2**. **(d)** Confocal images showing precise spatiotemporal control of OptoMT labeling of MTs within the regions indicated upon blue light exposure (470 nm, 40 μW/mm²). Also see **Supplementary Video 3**. **(e)** Confocal images showing robust and reversible OptoMT labeling of MTs across two successive dark-light cycles. Also see **Supplementary Figure 1**. **(f)** Confocal images of HeLa cells expressing mCh-OptoMT (red) co-stained with anti-α-tubulin (green) and DAPI (blue). Cells were either kept in the dark (top) or illuminated with blue light (470 nm, 40 μW/mm^2^, 30 sec) before fixation and immunostaining. **(g)** Live-cell imaging of HeLa cells co-expressing GFP-OptoMT (green in the merged panel) and H2B-mCh (red), showing MT cytoskeleton and mitotic progression at different cell-cycle stages before and after blue light exposure (470 nm, 40 μW/mm^2^, 5 sec). **(h)** Confocal images of L3 stage *C. elegans* expressing GFP::α-tubulin and mCh::OptoMT in epithelia. Blue light induces MT binding by mCh::OptoMT. (**i**-**j**) Confocal images (i) and quantitative analysis (j) of OptoMT-mediated reversible MT labeling in *C. elegans*. See **Supplementary Video 4**. The acquired data points were fitted by a single exponential decay function (t_1/2, on_ = 12.4 ± 3.2 sec; t_1/2, off_ = 252 ± 31 sec).

We next applied OptoMT to examine MT organization during mitosis. Using H2B as a chromatin marker, light-induced labeling enabled clear visualization of MT dynamics across distinct mitotic stages in live HeLa cells (**Fig. 1g**), without detectable perturbation on cell cycle progression, mitotic division, or viability (**Supplementary Fig. 2a-c**). To extend its application in vivo, we co-expressed GFP::tubulin (eGFP::TBB-2 β-tubulin) and mCherry::OptoMT under an epithelial-specific promoter in L3-stage *C. elegans*. GFP::tubulin revealed circumferential MT bundles in the syncytial epithelium, oriented perpendicular to the body axis (**Fig. 1h**). In the dark, mCherry::OptoMT was diffusely distributed throughout the epidermis, whereas blue light illumination triggered its rapid redistribution into patterned structures that closely colocalized with GFP::tubulin, confirming light-dependent MT labeling in vivo. OptoMT also exhibited robust and reversible MT labeling in *C. elegans*, with an activation half-life (t_1/2_, ON) of 12 s and a decay half-life (t_1/2_, _OFF_) of 252 s following light withdrawal (**Fig. 1i-j** and **Supplementary Video 4**). Thus, OptoMT recognizes *C. elegans* as well as human MTs.

Together, these results establish OptoMT as a single-component optogenetic probe that enables reversible and light-controlled MT labeling with minimal cellular perturbation, applicable both in vitro and in vivo.

### Design of photoswitchable MT plus-end trackers

To achieve reversible tracking of growing MT plus-ends, we adapted the optogenetic clustering strategy to MT tip-localization sequences containing the S/T-x-I-P (SxIP) consensus motif [1, 41, 42]. CRY2 was fused to SxIP-containing fragments of varying lengths derived from three representative MT plus-end tracking proteins (+TIPs), adenomatous polyposis coli (APC), dystonin (DST), and the stromal interaction molecule 1 (STIM1), which engage the end-binding protein 1 (EB1) at growing MT ends (**Fig. 2a-b** and **Supplementary Fig. 3a**) [37]. We reasoned that light-induced oligomerization of SxIP motifs could mimic their native multivalent assembly within scaffolding complexes, thereby enhancing EB1 recruitment and tip-tracking efficiency. Among the tested variants, CRY2-DST_5469-5485_ (V2, hereafter designated “OptoTIP”) showed the most robust comet-like localization upon blue light illumination (**Fig. 2c** and **Supplementary Fig. 3b-c**). OptoTIP displayed fast activation (t_₁/₂, ON_ = 11 s) and moderate deactivation (t_₁/₂, OFF_ = 204 s) kinetics, which were fully reversible across repeated dark-light cycles (**Fig. 2c**, **Supplementary Table 1**, and **Supplementary Video 5**). When co-expressed with GFP-EB1, mCherry-OptoTIP precisely colocalized with EB1 comets in a blue light-dependent manner, showing nearly identical fluorescence intensity profiles along MT tips (**Fig. 2d**, **Supplementary Fig. 3d**, and **Supplementary Video 6**). Nocodazole treatment, which depolymerizes MTs [43, 44], rapidly abolished the comet-like pattern formation even under continuous illumination, indicating that OptoTIP labeling is strictly dependent on intact MT polymer assembly (**Fig. 2c)**. No significant differences were observed in EB1 comet velocities between mCherry-(control) and mCherry-OptoTIP-expressing cells before and after blue light exposure, indicating that OptoTIP does not perturb endogenous MT growth (**Supplementary Fig. 3e**). Localized blue light stimulation within defined subcellular regions further induced spatially confined comet-like pattern formation, demonstrating the high spatiotemporal precision of this system (**Fig. 2e**).

**Figure 2.**
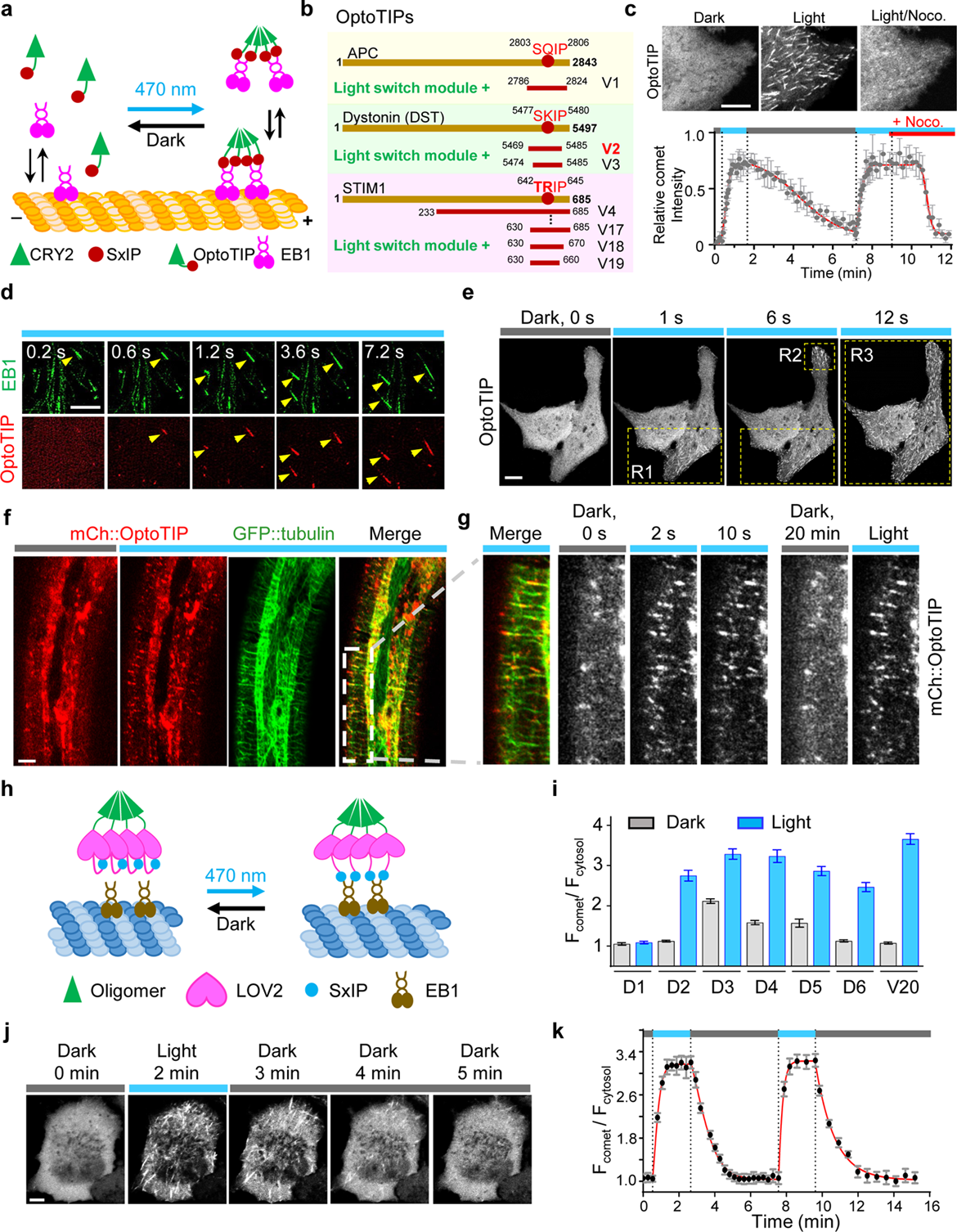
OptoTIP as a single component optogenetic actuator for reversible tracking of microtubule plus-ends (+TIPs). Scale bar, 5 μm. Error bars denote sem. **(a)** Schematic illustrating the design and light-induced association of OptoTIP with EB1 to track MT plus-ends. **(b)** OptoTIP variants were constructed by fusing CRY2 with EB1-binding motifs derived from APC, DST, or STIM1. **(c)** Confocal images and quantification of OptoTIP-mediated MT plus-end tracking during repeated dark-light cycles, followed by nocodazole treatment (20 μM). Blue bars indicate periods of illumination (470 nm, 40 μW/mm^2^). Data were fitted with a single exponential decay function (t_1/2, on_ = 11.1 ± 4.7 sec; t_1/2,off_ = 204 ± 44 sec). n = 10 cells from three independent biological replicates. Also see **Supplementary Video 5**. **(d)** Confocal images showing photo-activatable colocalization of mCh-OptoTIP (red) with GFP-EB1 (green) at growing MT plus-ends following blue light exposure, demonstrating faithful tracking of EB1-marked comets. Also see **Supplementary Video 6.** **(e)** Confocal images showing precise spatiotemporal control of OptoTIP tracking by visualization of successively selected photo-stimulation regions. **(f)** Confocal images of L3 stage *C.elegans* expressing GFP::α-tubulin and mCh::OptoTIP, showing blue light-inducible OptoTIP tracking of growing MT plus-ends. Also see **Supplementary Video 7.** **(g)** Enlarged region (white boxed inset) from panel (f) showing reversible OptoTIP relocalization upon alternating dark-light cycles.. Also see **Supplementary Video 8.** **(h)** Schematic depicting the design of LOV2-based OptoTIP variants. To enable tracking of MT plus-ends with faster deactivation kinetics, the SxIP motif derived from dystonin (DST_5469-5485_) was positioned downstream of the C-terminal Jα helix of LOV2, which was then fused either to a tetrameric DsRed (D series; D1-D6) or the light-inducible oligomerization module CRY2 (V20). **(i)** Quantification of comet-to-cytosol fluorescence intensity ratios for LOV2-based OptoTIP constructs. mCherry fluorescence at comets and adjacent cytosolic regions was measured before and after 30 s blue light exposure (470 nm, 40 μW/mm²). n = 17-25 cells from three independent biological replicates. **(j)** Confocal images of HeLa cells expressing mCherry-CRY2-LOV2-SxIP (V20, DST_5470-5485_) showing light-dependent plus-end tracking with rapid deactivation kinetics during sequential dark-light-dark cycles. Also see **Supplementary Video 9.** **(k)** Quantitative analysis of comet-to-cytosol fluorescence intensity ratios for the mCherry-CRY2-LOV2-SxIP (V20) construct. The measured kinetics revealed efficient MT plus-end tracking with an activation half-life of 9.1 ± 4.1 s and deactivation half-life of 40.2 ± 6.6 s. Data were fitted with a single exponential decay function. n = 16 cells from three independent biological replicates.

OptoTIP was robustly expressed across diverse cell types, including a dozen of cell lines derived from both excitable and non-excitable tissues, enabling light-inducible tracking of MT plus-ends in multiple cellular contexts (**Supplementary Fig. 3f**). To evaluate its performance in vivo, we co-expressed GFP::tubulin and mCherry::OptoTIP in the epidermis of *C. elegans* larvae, as above. Blue light illumination triggered robust and reversible tracking along GFP-labeled MT bundles on both dorsal and ventral epidermal surfaces (**Fig. 2f-g** and **Supplementary Videos 7-8**). Together, these results establish OptoTIP as a single-component optogenetic probe that enables light-inducible tracking of growing MT plus-ends at real time in living cells and animals without appreciable perturbation of endogenous cytoskeletal dynamics.

To expand the kinetic range of OptoTIP, we introduced photocycle mutations in CRY2 at residues L348 and W349 [45], located near the FAD binding site (**Supplementary Fig. 4a**). The L348F substitution produced a long-lived variant (t_1/2_, _OFF_ = 15.9 min), whereas W349 substitutions (e.g., W349H, W349L, W349E) either accelerated decay or abolished tip-tracking activity (**Supplementary Fig. 4b**). Nonetheless, all CRY2-based OptoTIP variants retained relatively slow recovery kinetics, with deactivation half-lives (t_1/2, OFF_) ranging from 2 to 16 min. To achieve faster deactivation, we adopted the LOV2 from oat phototropin, a blue light photosensory domain with a shorter photocycle [9, 10, 25, 46–48]. We placed the SxIP motif immediately downstream of the C-terminal Jα helix of LOV2, reasoning that EB1 binding would be sterically blocked in the dark and unmasked upon blue light-induced unfolding of the Jα helix (**Fig. 2h** and **Supplementary Fig. 4c**). We further fused LOV2-SxIP constructs to either DsRed (a constitutive tetramer) or CRY2 (a light-induced oligomer) to enhance avidity. After optimizing linker regions and the SxIP motif length (**Fig. 2i** and **Supplementary Fig. 4d**), we identified a LOV2-based OptoTIP variant (V20, **Supplementary Fig. 4d**) that exhibited rapid and reversible cytosol-to-comet transitions under repeated dark-ight cycles (**Fig. 2j-k**). This construct substantially accelerated deactivation kinetics, reducing the decay half-lives from minutes to approximately 40 s (**Fig. 2k**, **Supplementary Table 1**, and **Supplementary Video 9**). Together, these efforts yielded a toolkit of CRY2- and LOV2-based OptoTIP variants that enable reversible and light-tunable tracking of growing MT plus-ends, with the kinetic profiles ranging from seconds to minutes.

### OptoTIP as a tool to dissect STIM1-EB1 interactions

STIM1 is an ER-resdient Ca^2+^ sensor protein that activates store-operated Ca^2+^ entry (SOCE) by coupling to ORAI channels at the endoplasmic reticulum-plasma membrane (ER-PM) junctions. Beyond this canonical role, STIM1 can behave as a +TIP by hitchhiking on growing MT ends through an EB1-binding Ser/X-Ile-Pro (SxIP)-like motif (TRIP, residues 642-645), whereas its distal polybasic (PB) domain (residues 672-685) targets the PM via electrostatic interactions to engage ORAI channels [49–53]. These two C-terminal features likely create a competitive partition between MT plus-end tracking and PM engagement that shapes SOCE timing and localization [54–56].

To systematically define sequence determinants of STIM1 +TIP behavior, we performed deep mutational scanning of the conserved EB1-binding TRIP motif in the mCh-CRY2-STIM1_630-685_ construct (OptoTIP-V17, **Supplementary Fig. 3a**), generating a library of 80 variants to probe individual residue contributions to EB1 recognition (**Fig. 3a-c**). Upon blue light stimulation in HeLa cells, these mutants exhibited varying degrees of comet-forming activity (**Supplementary Fig. 5**). Positions I644 and P645 showed the highest stringency, as only Ile/Leu and Pro/Ala substitutions were tolerated, implying their key role in EB1 docking, whereas T642 and R643 were more permissive (**Fig. 3c** and **Supplementary Fig. 5**).

**Figure 3.**
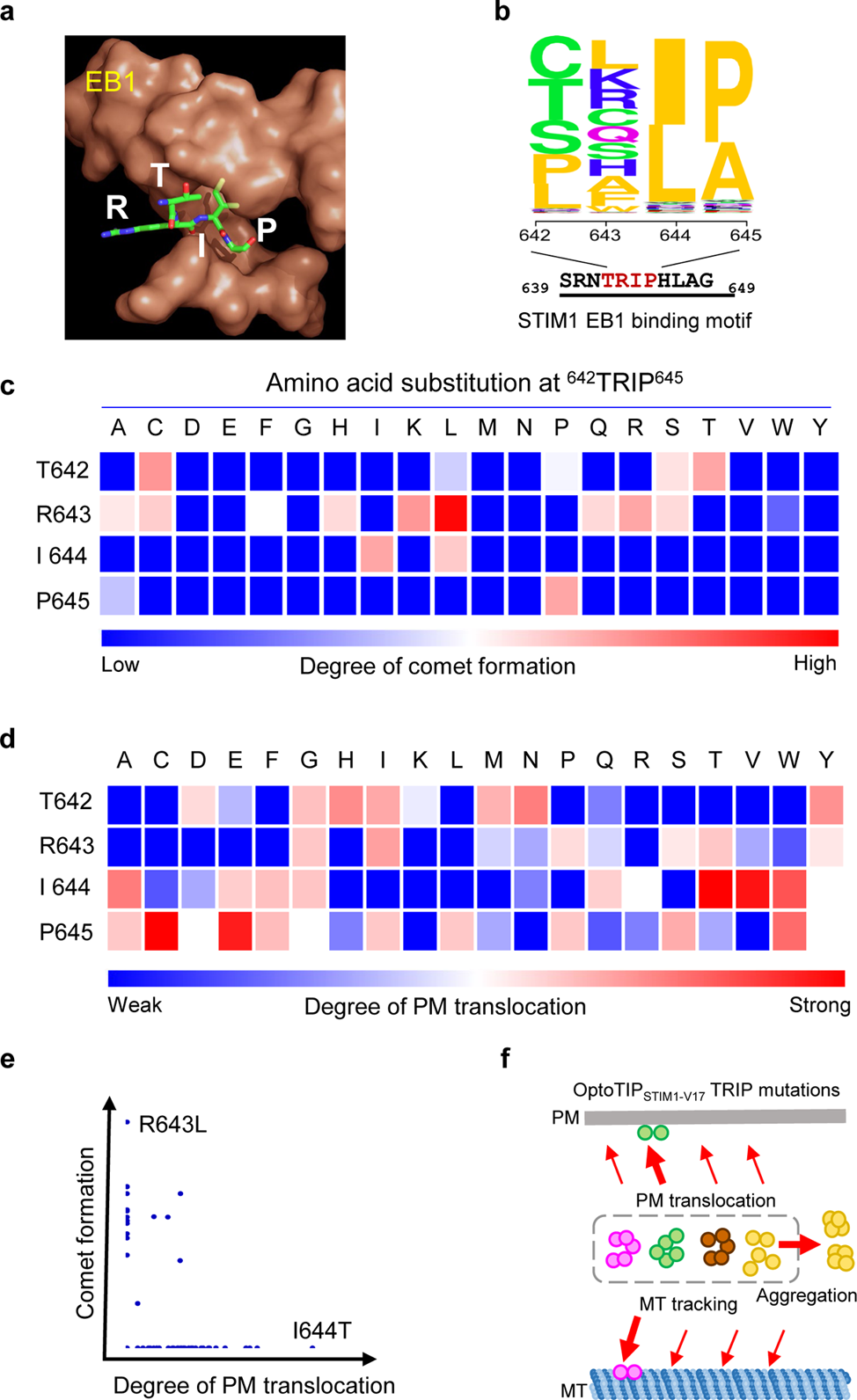
OptoTIP as an optogenetic probe to dissect determinatns governing STIM1-EB1 interactions. **(a)** Modeled 3D structure of the STIM1 EB1-binding motif (_642_TRIP_645_; shown as sticks) in complex with human EB1 (brown surface; PDB entry: 3GJO). **(b)** Sequence-logo representation of the tested EB1-binding motif derived from screening of 80 STIM1-TRIP mutants. The relative height of each residue reflects its MT plus-end tracking capability. OptoTIP-V17 construct was used in the assay (see **Supplementary Fig. 3a**), which was generated by fusing CRY2 to a fragment of the STIM1 cytoplasmic domain (aa 630–685). This region contains the EB1-binding TRIP motif (aa 642–645) and the polybasid domain (PB, aa 666–685) in the C-tail that mediates phosphoinositide interactions with the inner leaflet of the plasma membrane (PM). Also see **Supplementary Fig. 3** for representative images. (**c-d**) Heat-map representation of the extent of comet formation (**c**) or PM translocation (**d**) for all 80 OptoTIP-V17 variants. Each residue within the TRIP template (left column) was individually substituted with the 19 other amino acids indicated above the matrices. **(e)** Scatter plot summarizing the behaviors of 80 mCh-OptoTIP-V17 variants. Single-residue substitutions within the TRIP motif produced a bimodal distribution: mutants that lost plus-end tracking frequently accumulated at the plasma membrane, whereas those with enhanced plus-end tracking showed diminished membrane association, supporting a competitive interplay between EB1- and PM-targeting determinants within the STIM1 C-terminus. **(f)** Schematic summarizing four possible photo-inducible scenarios visualized in HeLa cells expressing mCh-OptoTIP-V17 variants: (i) MT plus-end tracking via EB1 binding; (ii) PM translocation via interactions with PM-resident phosphoinositides; (iii) diffuse cytosolic distribution; or (iv) aggregation due to CRY2 homo-oligomerization. Loss or weakening of EB1 binding typically results in one of the latter three outcomes.

We observed a clear reciprocal relationship across the mutant set: variants exhibiting strong MT comets seldom accumulated at the PM, whereas PM-enriched mutants showed diminished +TIP activity (**Fig. 3c-e** and **Supplementary Fig. 5**). This inverse partitioning supports the EB1-mediated diffusion-trap model, in which EB1 captures STIM1 at MT plus-ends and transiently restrains its ER-PM translocation [57]. Conversely, reduced EB1 affinity releases STIM1 to engage its C-terminal polybasic domain, facilitating PM anchoring and subsequent ORAI1 coupling. Congruently, these findings suggest that the native TRIP motif has evolved to confer an ideal affinity for EB1, which is strong enough to enable MT-plus-end surveillance yet sufficiently weak to permit timely PM engagement. Thus, EB1-TRIP and PB-PM interactions operate as opposing molecular levers, and OptoTIP provides a tunable platform to visualize and reprogram this finely balanced partitioning that governs SOCE dynamics (**Fig. 3f**).

### Optogenetic manipulation of tubulin post-translational modifications

We next explored whether OptoTIP or OptoMT could serve as modular platforms for light-inducible posttranslational modifications (PTMs) of tubulin. As a proof of concept, we focused on α-tubulin acetyltransferase (αTAT1), which catalyzes the acetylation of α-tubulin at the position K40 to enhance MT resistance to mechanical and chemical stress [58]. To render αTAT1 light-responsive, we incorporated the LOV2-based light-inducible nuclear export system (LEXY) [59] and two nuclear localization signals (NLS), yielding Opto-αTAT1. In the dark, NLS sequences confined the enzyme within the nucleus, whereas blue light triggered LEXY-mediated export to the cytoplasm (**Fig. 4a** and **Supplementary Fig. 6a**). For MT-specific recruitment, the N-terminal domain of CIB1 (CIBN, residues 1–81), which heterodimerizes with CRY2 under photostimulation [45, 60, 61], was appended to Opto-αTAT1 (**Supplementary Fig. 6a**). Upon blue light illumination, a portion of nuclear Opto-αTAT1 translocated to the cytosol (t_₁/₂, ON_ = 1.9 min; t_₁/₂, OFF_ = 1.7 min) and was subsequenlty recruited to OptoTIP- or OptoMT-decorated MT filaments through CRY2-CIBN heterodimerization (**Fig. 4b** and **Supplementary Fig. 6b-c**, and **Supplementary Video 10**). This relocalization induced a robust increase in tubulin acetylation, evidenced by enhanced acetyl-α-tubulin immunostaining (**Fig. 4c-d**) and progressive appearance of acetylated bands in Western blots after illumination (**Fig. 4e**). In contrast, control cells lacking Opto-αTAT1 only displayed basal acetylation levels (**Fig. 4d**).

**Figure 4.**
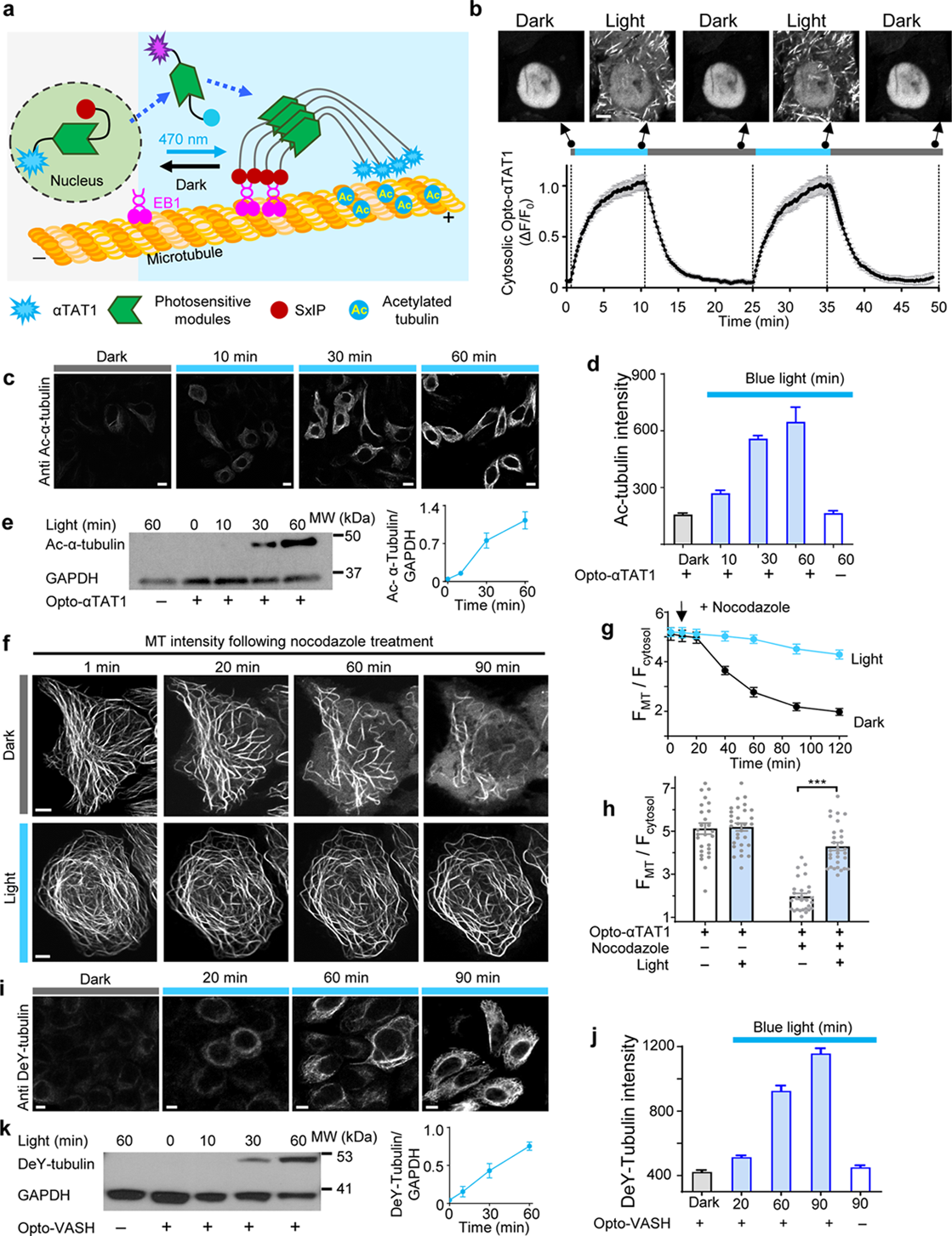
OptoMT or OptoTIP enables light-inducible posttranslational modifications of tubulin. Scale bar, 5 μm. Error bars denote sem. **(a)** A simplified schematic depicting the design of photosensitive acetyltransferase αTAT1, Opto-αTAT1. Upon blue light stimulation, Opto-αTAT1 undergoes nucleus-to-cytosol translocation, and is further recruited toward MT via light-dependent interaction with OptoTIP or OptoMT to efficiently catalyze the acetylation of α-tubulin. See **Supplementary Fig. 6** for details of the photosensitive modules. **(b)** Light-inducible nuclear export of mCh-Opto-αTAT1 with subsequent recruitment to MT in HeLa cells coexpressing GFP-OptoTIP. *Upper panel*, confocal images of the same cell before and after exposure to blue light at 470 nm (40 μW/mm^2^; blue bar). *Lower* panel, normalized cytosolic mCherry signals of Opto-αTAT1 during two repeated light-dark cycles. n = 24 cells from three independent experiments. Also see **Supplementary Video 10**. **(c)** Confocal images of HeLa cells co-expressing Opto-αTAT1 and OptoTIP after varying durations of blue light exposure (470 nm, 40 μW mm⁻²), followed by fixation and immunostaining for acetylated α-tubulin. **(d)** Quantification of α-tubulin acetylation levels from images in panel (c). Data represent 102 cells per time point from three independent biological replicates. **(e)** Immunoblot analysis of light-triggered α-tubulin acetylation (Ac–α-tubulin) in HeLa cells expressing Opto-αTAT1. Right, densitometric quantification showing the increased Ac–α-tubulin/GAPDH ratio upon blue light illumination. n = 3 independent biological replicartes. **(f)** Time-lapse confocal imaging of MT cytoskeleton (probed by GFP-OligoMT) in HeLa cells coexpressing OptoMT and Opto-αTAT1 in the presence of 2 μM nocodazole. Top panel (gray bar; dark), cells were cultured in the dark. For acquiring GFP signals, cells were very briefly subjected to photostimulation for 1-2 sec without eliciting nuclear export of Opto-αTAT1. Bottom panel (blue bar; light), cells were exposed to blue light (470 nm, 40 μW/mm^2^) before imaging with a pulse of 3 sec on and 30 sec off. **(g)** Quantification of the MT-over-cytosol fluorescence intensity ratio over 120 min of nocodazole treatment. n = 28 (dark) and 30 (light) cells from three independent biological replicates. **(h)** The bar graph and scatter dots showed the averaged values and distribution of MT-to-cytosol intensity ratios of GFP in cells cultured in the dark or under blue light illumination before and after nocodazole treatment. n = 28 (dark) and 30 (light) cells from three independent biological replicates. Each symbol represents the average of 5-6 measurements from one cell. \*\*\**P* < 0.001 (two-tailed Student’s *t*-test). **(i-k)** Light-dependent tubulin detyrosination enabled by Opto-VASH. **(i)** Confocal images of HeLa cells co-expressing Opto-VASH and OptoTIP after varying durations of blue light exposure (470 nm, 40 μW/mm^2^), followed by immunostaining for detyrosinated tubulin (DeY-tubulin). **(j)** Quantification of DeY-tubulin levels from images shown in panel (i). Data represent 96 cells per time point from three independent biological replicates. **(k)** Immunoblot analysis of DeY-tubulin in HeLa cells co-transfected with Opto-VASH and OptoTIP with or without blue light stimulation. Right, densitometric quantification of DeY-tubulin intensity. n = 3 independent biologial replicates.

To determine whether light-induced acetylation enhances MT stability, HeLa cells co-expressing GFP-OptoMT and Opto-αTAT1 were treated with nocodazole under illuminated or dark conditions. In the absence of light, nocodazole gradually disassembled the MT network within 90 min (**Fig. 4f-h**). Photoactivation of Opto-αTAT1, however, markedly increased MT resistance to depolymerization, largely preserving cytoskeletal integrity over the same period (**Fig. 4f-h**). These results demonstrate that Opto-αTAT1 enables light-dependent α-tubulin acetylation and strengthens the MT network against chemical stress.

Building on this approach, we next engineered a light-inducible detyrosination system, termed Opto-VASH, by fusing vasohibin 1 (VASH1), a tubulin carboxypeptidase that removes the C-terminal tyrosine from α-tubulin [62], to the same photosensitive module. When coupled with OptoMT, Opto-VASH progressively increased detyrosinated tubulin levels under pulsed blue light stimulation (3 sec ON, 30 sec OFF), as confirmed by both immunostaining (**Fig. 4i-j**) and Western blotting (**Fig. 4k**).

Together, these findings establish OptoTIP and OptoMT as modular scaffolds for temporal control of tubulin PTMs, which enables light-driven modulation of acetylation and detyrosination within the MT network of live cells.

### Optogenetic reconstruction of intracellular transport

Kinesin typically consists of an N-terminal motor domain, a central coiled-coil stalk that mediates dimerization and processive movement, and a C-terminal tail domain responsible for cargo binding and specificity [22, 63] (**Fig. 5a**, and **Supplementary Fig. 7a**). To render kinesin activity light-responsive, we replaced the native coiled-coil stalk and C-terminal tail with CRY2 (**Fig. 5b**), reasoning that blue light-induced CRY2 oligomerization could reconstitute motor dimerization and activation (**Fig. 5c**).

**Figure 5.**
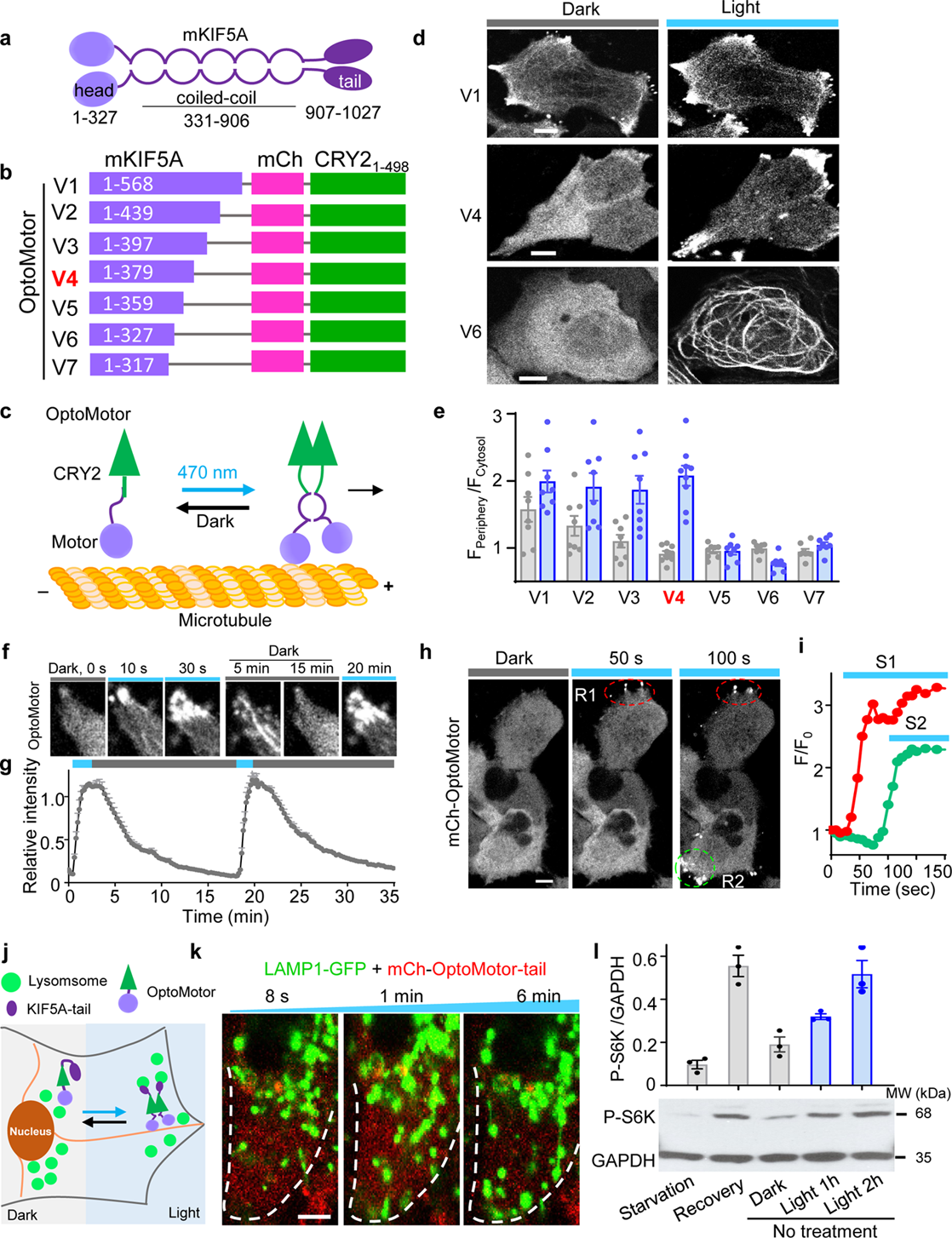
OptoMotor designed to photo-manipulate intracellular transport. Scale bar, 5 μm. Error bars denote sem. **(a)** Domain organization of mouse KIF5A (mKIF5A), showing the N-terminal motor head, central coiled-coil stalk, and C-terminal cargo-binding tail. **(b)** Schematic of mKIF5A truncation variants fused to CRY2 for optogenetic activation. Variant V4 (residues 1-379) was identified as the optimal construct and designated OptoMotor. **(c)** Design principle of OptoMotor. Blue light illumination induces CRY2 oligomerization to drive reassembly of the truncated motor complex and restoring plus-end-directed motility along under MTs. **(d)** Confocal images of selected CRY2-KIF5A truncation variants expressed in HeLa cells under dark and illuminated conditions. Variant V1 exhibited constitutive peripheral accumulation; V4 showed robust light-induced redistribution to the cell periphery without appreciable basal activity; and V6 displayed pronounced MT labeling. See **Supplementary Fig. 7** for the complete set of variants. **(e)** Quantification of periphery-to-cytosol fluorescence ratios across truncation variants. Each symbol represents the mean of 5-6 cells from a single imaging field. A total of 40-50 cells were analyzed per construct across three independent biological replicates. **(f)** Time-lapse images showing reversible distribution of OptoMotor during repeated dark-light cycles. Also see **Supplementary Video 11**. **(g)** Kinetic trace of peripheral intensity changes showing OptoMotor activation (t_1/2_, _ON_ = 0.9 min) and deactivation (t_1/2_, _OFF_ = 3.7 min). **(h)** Localized blue light illumination within defined regions of interest (ROIs R1, R2) elicited spatially confined redistribution of OptoMotor in single cells. **(i)** Quantification of relative fluorescence intensity changes within ROIs confirming high spatial precision of light-triggered activation. **(j)** Schematic of the chimeric OptoMotor-Tail construct, generated by fusing the cargo-binding C-terminal tail of mKIF5A (residues 907-1027) to OptoMotor. Upon blue light activation, OptoMotor-Tail drives peripheral transport of lysosomes to enhance mTORC signaling. **(k)** Confocal images of HeLa cells co-expressing LAMP1-GFP and mCh-OptoMotor-Tail showing light-induced redistribution of lysosomes toward the cell periphery. **(l)** Immunoblot showing phosphorylation of S6K at T389 (P-S6K) as a readout of mTORC1 activity. Blue light illumination enhanced P-S6K levels in cells expressing OptoMotor-Tail, consistent with lysosomal repositioning mediated activation of mTORC1 signaling. Starved cells before and after nutrient recovery were used as negative and positive controls for reporting mTORC1 activity.

Using KIF5A, a kinesin-1 heavy chain with well-characterized biophysical properties [64, 65], as the engineering scaffold, we systematically truncated its coiled-coil region to generate seven CRY2-fused variants with progressively shortened neck linkers (**Fig. 5b**). This series enabled identification of configurations that preserved processive stepping while supporting blue light-dependent activation (**Fig. 5b-c**). When expressed in HeLa cells, the variants displayed distinct localization patterns before and after light illumination.

Constructs retaining longer coiled-coil regions (beyond residue 397; V1-V3) accumulated at the cell periphery even in the dark, suggesting that excessive coiled-coil inclusion promotes spontaneous plus-end targeting (**Fig. 5d-e**, and **Supplementary Fig. 7b**). The shorter truncations behaved differently, with KIF5A_1-327_-CRY2 (V6) remained cytoplasmic in the dark yet binding strongly to MTs upon activation (**Fig. 5d**), functioning as a light-inducible MT-binding probe. By contrast, KIF5A_1-359_-CRY2 (V5) remained unresponsive to light, while KIF5A(1-317)-CRY2 (V7), which harbors an incomplete motor domain, formed punctate aggregates likely arising from intrinsic CRY2 clustering (**Fig. 5e**, and **Supplementary Fig. 7b**). Among these, KIF5A_1-379_-CRY2 (V4, termed OptoMotor) showed optimal performance, remaining diffusely cytoplasmic in the dark and rapidly redistributing to the cell periphery under blue light (**Fig. 5d-e**, and **Supplementary Video 11**). Quantitative analysis revealed rapid and reversible light-inducible activation, with an activation half-life (t_₁/₂, ON_) of 0.9 min and a deactivation half-life (t_₁/₂, OFF_) of 3.7 min (**Fig. 5f**-**g**, and **Supplementary Video 11**). Localized blue light stimulation further induced region-specific redistribution of OptoMotor, confirming its high spatiotemporal precision and controllability (**Fig. 5h-i**). Together, these results establish OptoMotor as a single-component light-inducible hybrid motor capable of reconstituting processive plus-end-directed transport.

Because the C-terminal tail domain of kinesin-1 governs cargo specificity and lysosomal transport [66, 67], we next examined whether OptoMotor can be coupled with naturally evolved tail domain to drive cargo movement and functional signaling in a light-dependent manner. Appending the native tail domain (residues 907-1027) to OptoMotor yielded OptoMotor-Tail (**Fig. 5j**). In HeLa cells co-expressing OptoMotor-Tail, the majority of LAMP1-GFP labeled lysosomes accumulated near the nucleus with minimal bidirectional movement in the dark, whereas blue light stimulation triggered their outward movement toward the cell periphery (**Fig. 5k**). This light-driven redistribution demonstrates that OptoMotor-Tail restores cargo recognition and motor-driven lysosomal transport in a controllable manner.

Peripheral redistribution of lysosomes promotes mTORC1 activation by positioning the organelles closer to Rheb-enriched membranes and growth factor-responsive signaling domains at the cell periphery [68–72]. Consistent with this, photoactivation of OptoMotor-Tail and the ensuing peripheral repositioning of lysosomes resulted in a marked increase in phosphorylation of ribosomal protein S6 kinase (S6K) at T389 (**Fig. 5l**), a hallmark readout of mTORC1 activation that reflects enhanced anabolic signaling and protein synthesis [73, 74]. Thus, light-induced recruitment and activation of OptoMotor-Tail directly promote lysosomal repositioning, which in turn boosts mTORC1 signaling through spatial coupling of the lysosomal nutrient-sensing machinery.

Together, these results establish OptoMotor as an optogenetic actuator that reconstitutes intracellular transport and functionally couples cytoskeletal dynamics to signaling output, providing compelling evidence that spatial control of lysosomal trafficking can causally modulate mTORC1 activity.

### Optogenetic engineering of spastin as a modular MT-severing tool

Having established optogenetic platforms for light-tunable labeling, plus-end tracking, post-translational modification, and cargo transport, we next sought to extend this framework to active remodeling of the MT cytoskeleton itself. While the preceding modules primarily enabled visualization or modulation of existing MT structures, a tool that could selectively disassemble MT filaments with spatiotemporal precision would complete the functional continuum of our toolkit: from construction and transport to controlled depolymerization and turnover.

Spastin, a hexameric AAA ATPase, catalyzes MT severing by extracting tubulin subunits in an ATP-dependent manner [27]. Its activity is orchestrated by multiple domains, including an N-terminal MIT domain that governs subcellular targeting, a microtubule-binding domain (MTBD) that recognizes tubulin C-terminal tails, and a C-terminal AAA domain that mediates ATP hydrolysis and filament severing [26, 75]. Because enzymatic activation requires hexamerization, we reasoned that light-induced oligomerization could be leveraged to reconstitute spastin activity in a reversible and tunable fashion.

Building on the OptoMT and OptoTIP scaffolds, we employed a truncation-guided strategy to develop a light-inducible MT-severing toolkit (**Fig. 6a**). Spastin truncations spanning residues 2-616, 228-616, 245-616, 270-616, and 343-616 were fused to OptoMT and expressed in HeLa cells, and the MT network integrity was assayed with GFP-OligoMT, a genetically encoded reporter we previously developed to visualize the MT cytoskeleton [76]. Among the tested constructs, spastin_228-616_ produced the most pronounced filament fragmentation and fluorescence loss after 2 hours bule light inllumination (**Supplementary Fig. 8a-b**). Because spastin_228-616_ retains the MTBD region, we speculated that residual MT engagement could confer background activity. To test this independent of OptoMT targeting, we expressed mCherry-CRY2-spastin_228-616_ and observed partial MT disruption upon blue light stimulation, evidenced by cytosolic diffusion of OligoMT signal with concominant reduction of filamentous structures (**Supplementary Fig. 8c**).

**Figure 6.**
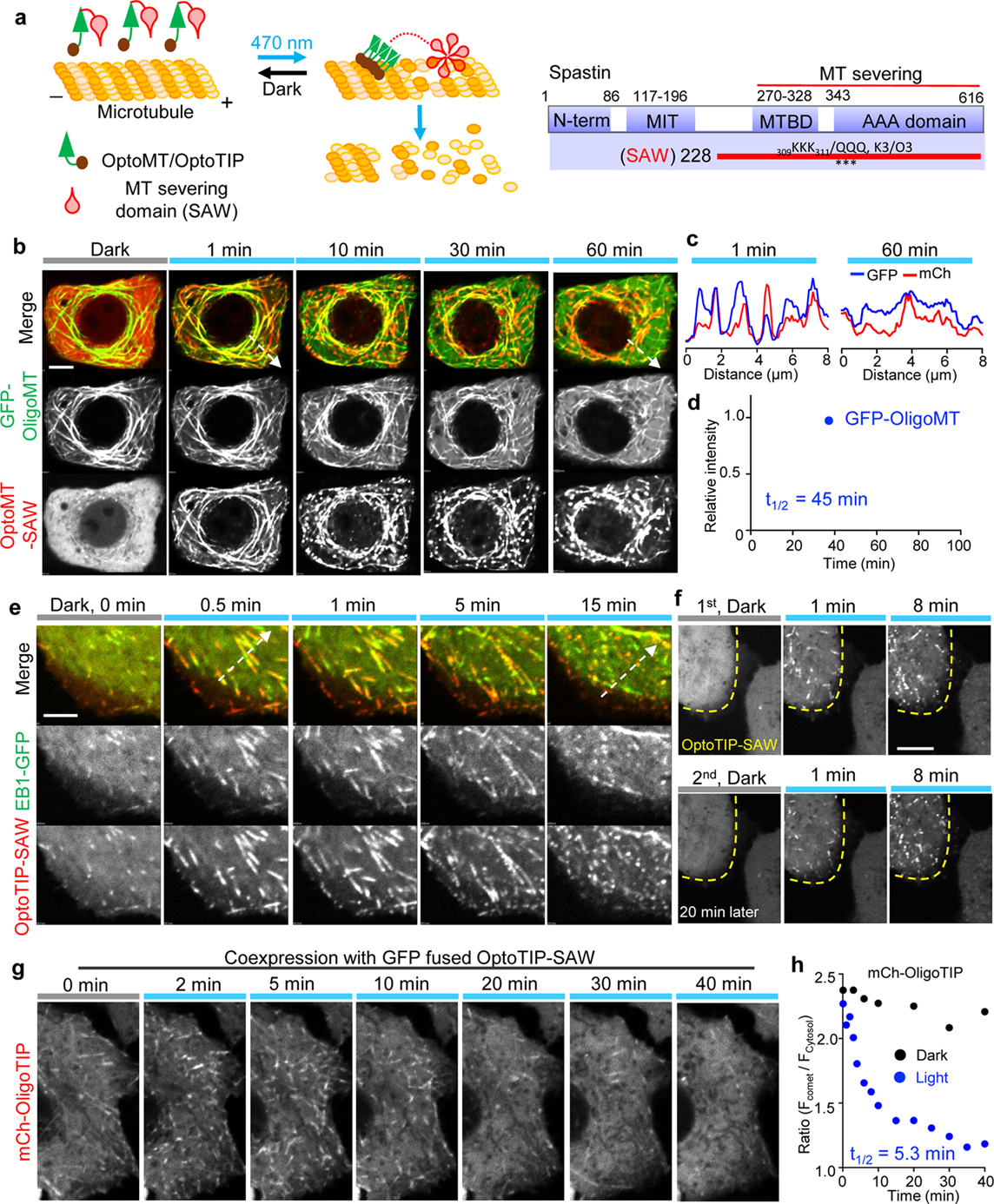
OptoSAW enables precise control of microtubule severing. Scale bar, 5 μm. Error bars denote sem. **(a)** Schematic illustration of the single-component OptoSAW constructs generated by directly fusing the minimal MT-severing domain of spastin (residues 228-616 carrying the K3/Q3 mutation, ³⁰⁹KKK³¹¹ → QQQ) to either OptoMT or OptoTIP. See **Supplementary Fig. 8** for spastin engineering and characterization details. **(b)** Time-lapse confocal images monitoring light-inducible MT severing in single HeLa cells expressing mCh-OptoMT-SAW (red) and GFP-OligoMT (green), the latter serving as a MT marker. See **Supplementary Video 12**. **(c)** Fluorescence intensity profiles of GFP (green) and mCherry (red) signals measured along the dashed line shown in panel b, demonstrating the progressive decline of filament continuity during blue light illumination. **(d)** Quantification of filament-associated GFP intensity from images shown in panel b. n = 32 cells from three independent biological replicates. **(e)** Time-lapse confocal images of HeLa cells co-expressing EB1-GFP (green) and mCherry-OptoTIP-SAW (red), showing light-induced disruption of +TIP comet dynamics. See **Supplementary Video 13**. **(f)** Confocal images demonstrating spatially restricted MT severing. Patterned blue light illumination selectively induced MT disassembly within targeted regions. **(g)** Time-lapse imaging of HeLa cells co-expressing mCherry-OligoTIP (+TIP marker) and OptoTIP-SAW following blue light stimulation, revealing progressive disassembly of +TIP comets. **(h)** Quantitative analysis of the ratio of the comet-to-cytosolic mCherry fluorescence intensities in cells expressing OptoTIP-SAW under dark (black) and illuminated (blue) conditions, demonstrating efficient, light-dependent +TIP severing. n = 16 cells from three independent biological replicates.

To eliminate this basal activity, we introduced charge-neutralized mutations (residues 309-311 KKK→ QQQ, K3/Q3) within the MT-binding domain, which minimized basal severing and preserved MT integrity (**Supplementary Fig. 8d**). When the catalytically competent but non-MT-binding spastin_228-616_ -K3/Q3 mutant (designated SAW, which stands for Severing Activated by Wavelength) was appended to either OptoMT or OptoTIP (designated OptoMT-SAW or OptoTIP-SAW, respectively), both chimeras triggered robust and spatially confined MT disassembly upon blue light exposure (**Supplementary Fig. 9a-e**). These results indicate that efficient severing requires both MT targeting and light-driven oligomerization. Upon activation, OptoMT-SAW induced progressive fragmentation and cytoplasmic dispersion of OligoMT-labeled filaments (**Fig. 6b-c** and **Supplementary Video 12**), with a severing half-life of approximately 45 min (**Fig. 6d**).

Likewise, OptoTIP-SAW efficiently severed MT plus-end structures, as indicated by the progressive loss of comet-like +TIPs visualized with EB1-GFP (**Fig. 6e**, and **Supplementary Video 12**). Under repeated light–dark cycles, blue light selectively recruited OptoTIP-SAW to defined ROIs, inducing localized severing of +TIPs while neighboring non-illuminated cells retained diffuse cytoplasmic signal. After 20 min in the dark, OptoTIP-SAW remained responsive, relabeling and severing +TIPs exclusively within the previously illuminated ROI (**Fig. 6f**). To quantify severing kinetics while avoiding potential artifacts from EB1-based labeling, we employed OligoTIP, a genetically encoded +TIP marker that does not interact with OptoTIP constructs [76]. Analysis of comet-to-cytosol fluorescence ratios revealed rapid loss of +TIP integrity, with a severing half-life of around 5.3 min (**Fig. 6g-h**, and **Supplementary Video 13**). These findings establish OptoTIP-SAW as a reversible, spatially programmable optogenetic module for precise disassembly of dynamic MT plus-end structures in living cells. Together, these results establish OptoMT-SAW and OptoTIP-SAW as single-component programmable optogenetic actuators (collectively termed OptoSAW) for targeted MT disassembly in living cells.

### OptoSAW applied to interrogate cell signalling and organelle transport

We next used OptoSAW to examine how acute disruption of dynamic MTs reshapes STIM1-medited cell signaling and organelle trafficking in living cells. While STIM1 primarily functions as an ER calcium sensor, its interaction with EB1 couples the ER to dynamic MT plus-ends, promoting membrane remodeling and modulating the timing of its translocation to ER-PM junctions and then regulating store-operated calcium entry (SOCE) (**Fig. 7a**). We firslty examined how MT disassembly influences calcium influx. Depletion of ER calcium store using thapsigargain (TG) elicited comparable calcium influx, as assessed by GCaMP6s fluorescence, in both control and OptoTIP-SAW expressing cells after 30 min of illumination (**Fig. 7b**). However, STIM1 puncta formed more rapidly at ER-PM junctions following MT disassembly (**Fig. 7c-d**, and **Supplementary Video 14**). This is consistent with the model in which EB1 binding tethers STIM1 to growing MT plus-ends, transiently delaying its translocation to the ER-PM junctions and preventing excessive calcium entry (**Fig. 7a**). Disruption of MT plus-end dynamics therefore accelerates STIM1 recruitment to ER-PM junctions without altering overall SOCE amplitude. Together, these results suggest that while MT are dispensable for the magnitude of SOCE, they probably fine-tune the temporal dynamics of STIM1 activation.

**Figure 7.**
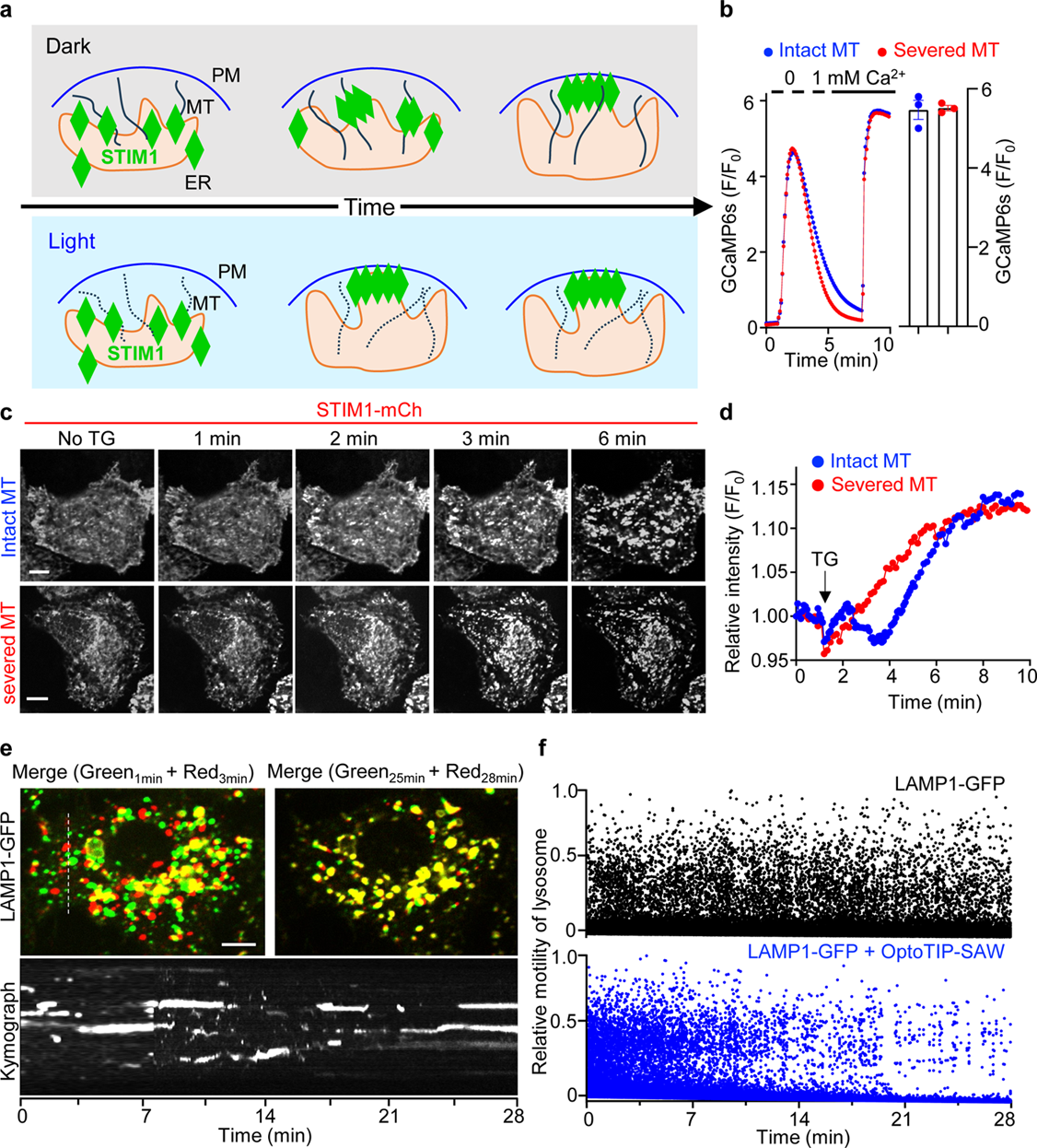
OptoTIP-SAW dissects microtubule-dependent signaling and organelle transport in real time. Scale bar, 5 μm. Error bars denote sem. **(a)** Schematic illustrating that disruption of MT plus-ends releases activated STIM1, promoting its translocation to ER-PM junctions and subsequent ER membrane remodeling. **(b)** Representative fluorescence traces (left) show Ca²⁺ influx in control cells (blue, intact MTs) and OptoTIP-SAW–expressing cells (red, MT severed). Bar graph on the right shows the quantification of SOCE amplitude from three independent biological replicates (Each dot represents the mean of 30–60 cells). Assays were performed in GCaMP6s-HeLa stable cells expressing mCherry-OptoTIP (non-severed control, blue) or mCherry-OptoTIP-SAW (red). Cells were illuminated for 30 min before store depletion with thapsigargin (1 μM). **(c-d)** Accelerated STIM1 activation kinetics following MT disassembly. (**c**) Confocal of HeLa cells stably expressing STIM1–mCherry and ORAI1–GFP, either without (blue, intact MTs) or with (red, MT severed) co-expression of miRFP-OptoTIP-SAW. Cells were illuminated for 2 h before ER store depletion with 1 μM TG, showing faster formation of STIM1 puncta in the MT-severed condition. (**d**) Quantification of STIM1 puncat formation kinetics following TG-induced store depletion. Also see **Supplementary Video 14**. **(e-f)** OptoTIP-SAW enables light-controlled disruption of lysosomal transport. (**e**) Merged confocal images showing the subcellular distribution of LAMP1-GFP labeled lysosomes at the indicated time points. Top left, merged images comparing 1 min (green) and 3 min (red); top right, 25 min (green) and 28 min (red). Bottom, kymograph along the dashed line (top left) illustrating progressive attenuation of lysosome motility over 28 min. (**f**) Quantification of motile LAMP1-GFP-labeled lysosomes in cells with or without OptoTIP-SAW expression. Also see **Supplementary Video 15**.

We next applied OptoSAW to investigate how dynamic MTs contribute to organelle trafficking, focusing on lysosomal transport as a representative example. Bidirectional lysosome transport along MT tracks is essential for cellular adaptation and homeostasis in response to environmental cues and stress [70, 77]. For instance, nutrient deprivation drives retrograde lysosome movement toward MT minus-ends near the centrosome via cytoplasmic dyneins, whereas nutrient repletion or growth factor stimulation promotes kinesin-mediated anterograde lysosome movement toward MT plus-ends at the cell periphery [68, 78]. We leveraged OptoTIP-SAW to assess how disruption of MT dynamics affects lysosomal trafficking. In control HeLa cells, LAMP1-GFP-labeld lysosomes exhibited canonical bidirectional, stop-and-go motion with dynamic redistribution over time (lower panel; **Supplementary Video 15**). In contrast, co-expression of Opto-TIP-SAW caused a progressive decline of lysosomal motility under blue light exposure, culminating in transport arrest within 30 min (**Fig. 7e**, and **Supplementary Video 15**). Kymographs analysis further revealed a shift from dispersed to stationary trajectories, confirming a time-dependent reduction in motile lysosomes (**Fig. 7f**). Collectively, these results demonstrate that light-induced disassembly of MT plus-ends by OptoTIP-SAW effectively impairs MT-based lysosomal transport, confirming the critical role of dynamic MTs in sustaining organelle motility.

## DISCUSSION

MTs form the dynamic scaffold that supports cellular organization, acting not only as structural elements but also platforms for force generation, intracellular cargo transport, organelle positioning, and signaling coordination [79–82]. Their highly adaptable nature, shaped by an intricate network of microtubule-associated proteins (MAPs), confers remarkable architectural plasticity while enabling rapid reorganization in response to cellular cues. Yet this very complexity has made it challenging to define how local MT dynamics translate into functional outcomes within living cells. Coventional approaches for labeling or perturbing MTs, such as pharmacological agents or genetic manipulation, lack the temporal control, reversibility, and subcellular precision needed to capture transient remodeling events. Here, we present a unified modular optogenetic framework that overcomes these barriers by enabling precise, reversible control of MT labeling, post-translational modification, motor activity, and filament severing. By converting light-induced clustering into tunable MT association, this single-component toolkit provides a versatile means to visualize, manipulate, and reconstitute MT behavior in living systems.

A central advance of this work lies in its minimalist, single-component design, which harnesses light-dependent clustering to program MT engagement. OptoMT and OptoTIP represent a new class of reversible, photoswitchable MT probes with minimal dark-state activity and subcellular precision. Compared with conventional visualization methods [83–85], these tools avoid artifacts arising from overexpression, fixation, or chemical labeling, and remain fully compatible with long-term live-cell imaging. Their activation and recovery kinetics, which is tunable from seconds to minutes, complement existing optogenetic strategies while offering higher temporal precision and flexibility. Notably, prolonged expression of OptoMT or OptoTIP did not affect cell viability, mitotic progression, or MT organization under either dark or illuminated conditions, indicating their non-perturbative nature. Although previous optogenetic designs have targeted EB1 or related MAPs[18, 86, 87], these efforts largely focused on partner recruitment or inhibition rather than on reversible labeling. Our system extends the optogenetic repertoire to enable fully reversible MT labeling and tracking within a single genetic module. Using STIM1 as a test case, we showed that OptoTIP allows systematic dissection of +TIP-MT interactions in living cells. This strategy is broadly adaptable for mapping other MAPs and decoding motif-dependent MT targeting with high spatial and temporal resolution.

Beyond visualization, the same framework supports localized biochemical editing of the MT lattice. Tubulin post-translational modifications (PTMs) critically influence filament stability, motor affinity, and intracellular organization [4]. By fusing α-tubulin acetyltransferase (αTAT1) or vasohibin-1 (VASH1) to our light-responsive scaffolds [58, 88], we achieved optical control of tubulin acetylation and detyrosination, respectively. This capability enables spatiotemporally resolved manipulation of local MT chemistry and reveals direct links between site-specific PTMs and filament stability. These findings complement efforts to decipher the “tubulin code” [89] and can be readily extended to other PTMs such as glutamylation or glycylation [90, 91]. The ability to impose defined chemical marks on demand provides a route to probe how local biochemical cues coordinate processes such as cell polarity, mitosis, and organelle trafficking.

To reconstruct motor-driven transport, we engineered OptoMotor, a minimal kinesin actuator in which the endogenous coiled-coil stalk is replaced by a light-inducible oligomerization domain. This design preserves the intrinsic gating mechanism of kinesin while allowing optical control of dimerization and motility. Fusion of the endogenous tail restored cargo specificity, enabling light-dependent redistribution of lysosomes. Remarkably, peripheral lysosome repositioning upon OptoMotor-Tail activation coincided with enhanced phosphorylation of S6K at T389, a canonical mTORC1 substrate [73, 74, 92], demonstrating a direct mechanistic link between lysosomal positioning and mTORC1 signaling. Unlike two-component recruitment systems that depend on artificial tethering or cargo overexpression [24, 93–95], OptoMotor operates as a self-contained module that maintains native topology, facilitating direct analysis of how motor assembly and cargo engagement shape compartmentalized signaling outputs.

For active remodeling of the cytoskeleton, we developed OptoSAW, a compact, spastin-derived actuator that achieves light-controlled MT disassembly through tunable oligomerization. Microtubule-severing enzymes such as spastin and katanin remodel the cytoskeleton by extracting tubulin subunits from the lattice [26, 27], yet previous optical systems for MT severing typically required multicomponent assemblies [16, 96]. Our single-component design simplifies genetic delivery, maintains balanced stoichiometry, and enables precise spatial control. Light-driven oligomerization of the spastin core allowed inducible MT severing. Acute disruption of dynamic plus-ends reduced lysosomal motility and altered ER architecture, clearly demonstrating the central role of MT integrity in coordinating organelle transport and signal transduction.

Several technical considerations warrant mention. Blue light activation limits penetration depth and can introduce phototoxicity, motivating future development of red-shifted or near-infrared photoreceptors to extend in vivo applicability [11, 12, 97]. The magnitude of CRY2 clustering depends on expression level and temperature, suggesting that standardized promoters or knock-in models will enhance reproducibility. Enzymatic modules such as αTAT1 may exceed physiological catalytic rates and thus require calibration to native activity. Sustained activation of OptoSAW can cause irreversible lattice loss, emphasizing the need for optimized illumination duty cycles. Finally, achieving multiplexed control across distinct wavelengths remains an engineering challenge; the addition of spectrally orthogonal switches or self-limiting degradation motifs could refine temporal precision and expand multicolor compatibility.

In summary, this single-component optogenetic platform bridges molecular engineering and cell physiology by integrating reversible visualization, local biochemical modification, motor regulation, and filament remodeling within a single photoregulatory framework. Through programmable control of MT dynamics, it transforms optogenetics from a localization-based perturbation tool into a quantitative system for reconstructing cytoskeletal logic in living cells. The ability to optically build, reshape, and dismantle MT networks in real time establishes a foundation for decoding how cytoskeletal architecture orchestrates intracellular signaling, organelle positioning, and disease-related remodeling.

## METHODS

### Reagents and antibodies

KOD Hot Start DNA Polymerase was purchased from EMD Millipore Corporation. Restriction endonucleases, NEBuilder HiFi DNA Assembly Master Mix and T4 DNA ligase were obtained from New England BioLabs. QuikChange Multi Site-directed Mutagenesis Kit was purchased from Agilent Technologies. Nocodazole (CAS, 31430-18-9) and thapsigargin (CAS, 67526-95-8) were purchased from Sigma-Aldrich. These compounds were dissolved in DMSO and prepared as stock solutions (1 mg/mL). The anti α-tubulin, acetyl-α-tubulin (Lys40) antibodies were purchased from Santa Cruz Biotechnology (Cat No. SC-32293 and SC-23950). Tubulin Tracker (Deep Red, Cat No. T34077) and Goat anti-mouse IgG highly cross-adsorbed secondary antibodies (conjugated with Alexa Fluor 488 or 555; Cat No. #A-11001, and A-21422) were purchased from Thermo Fisher Scientific.

### Plasmid construction

The plasmid templates for EB1 (#17234), CLIP170 (#54044), CAMSAP1 (# 59036),CAMSAP2 (#59037), KIF5A (#166954), and spastin (#134461) were purchased from Addgene. To generate OptoMT, we firstly amplified the PHR domain (residues 1-498) of *Arabidopsis thaliana* CRY2 (Addgene, #70159) by standard PCR and then inserted the fragment into modified pmCherry-C1 and pEGFP-C1 vectors (Clontech), followed by the insertion of multiple MT binding domains, derived from EB1, CLIP170, KIF5A, or CAMSAP1 at the BspEI and EcoRI/BamHI sties, as well as a 3x(SGGGGG) flexible linker between CRY2 and the MT binding domain. For OptoTIP, the EB1-binding SxIP motifs were either directly synthesized, derived from DST (residues 5469-5485), DST (residues 5474-5485), or APC (residues 2786-2824), by Integrated DNA Technologies or amplified via standard PCR. These fragments were inserted to replace the MT binding domain within OptoMT to generated mCh-OptoTIP or EGFP-OptoTIP. OptoTIP variants were subsequently made using the QuikChange multi site-directed mutagenesis Kit (Agilent). To produce LOV2-based OptoTIP, LOV2 from oat phototropin was amplified and then inserted between CRY2 and the SxIP motif. The linker and LOV2 junction regions were further optimized by standard PCR. Opto-αTAT1 (αTAT1-mCh-CIBN-NLS-LEXY) was constructed by using the HiFi DNA assembly method. αTAT1, mCherry, CIBN and LEXY (LOV2-NES) were amplified by standard PCR from pEF5B-FRT-GFP-αTAT1 (Addgene, #27099), pmCherry-C1, CIB1-CreC(N1) (Addgene, #75367) and NLS-mCherry-LEXY (Addgene, #72655), respectively. The amplified fragments were ligated by using the NEBuilder HiFi DNA assembly enzyme (New England Biolabs).

To generate OptoMotor constructs, the sequences of motor-contained domains from kinesin-1 (mouse KIF5A, Addgene, #166954 and mouse KIF5B, Addgene, #31604), kinesin-3 (mouse KIF1B, DNAsu, #MmCD00083530 and human KIF1C, DNAsu, #HsCD00438806), and kinesin-4 (human KIF4B, DNAsu, #HsCD00295076) were amplified by PCR and inserted upstream of CRY2 in either the pEGFP-N1 or pmCherry-N1 vector. To generate OptoMotor constructs with cargo-binding specificity, the C-terminal tail of KIF5A (residues 803-1027) was PCR-amplified and inserted downstream of the CRY2 domain using BspEI and BamHI sites. Organelle-specific targeting was achieved by co-expressing CIBN fused to mitochondrial (AKAP1) or lysosomal (LAMP1) membrane anchors.

To construct the OptoSAW constructs, truncated MT-severing domains of human spastin (Addgene, #134461) were PCR-amplified and fused to the C-terminus of OptoMT or OptoTIP. To minimize background MT binding of spastin, a triple-lysine motif (K310-K312) critical for MT association was mutated to glutamines (KKK-QQQ) using the QuikChange multi site-directed mutagenesis kit (Agilent). All plasmids were verified by Sanger sequencing prior to downstream applications.

### Cell culture and transfection

HeLa and other cell lines, including HEK293, SK-MEL-28, C2C12, U87, NIH3T3, MIA PaCa-2, COS-7, MEF, Neuro 2A, H9C2, and A549, were obtained from ATCC and maintained at 37 °C with 5% CO2 in complete cell-culture medium according to recommendations from the supplier. DNA transfection was carried out by using Lipofectamine 3000 (Life Technologies) following the manufacturer’s instructions. For live-cell or fixed-cell imaging experiments, cells were seeded in four-chamber 35-mm glass-bottom dishes (D35C4-20-1.5-N, Cellvis) at 20-40% confluency one day before transfection.

### Immunostaining

HeLa Cells were seeded on four-chamber 35-mm glass-bottom dishes and cultured until reaching 60-80% confluency. Cells were then transfected with the indicated plasmids using Lipofectamine 3000 following the manufacturer’s instructions. 16-24 h post-transfection, cells were either kept in the dark or exposed to blue light illumination (470 nm, 40 μW/mm^2^) with the indicated pulse and duration. The cells were fixed with 4% paraformaldehyde in PBS for 20 min at room temperature (RT), rinsed three times with PBS, and permeabilized with 0.1% Triton X-100 in PBS for 10 min at RT. Fixed cells were then blocked with 10% goat serum (Thermo Fisher, #50062Z) for 1 h at RT and incubated with a rabbit anti-α-tubulin antibody (1:200 dilution) overnight at 4 °C. After three washes with PBST, cells were incubated with Alexa Fluor 488- conjugated IgG secondary antibody (1:500 dilution) for 1 h at RT. Nuclei were counterstained with DAPI, and samples were washed thoroughly and stored in PBS prior to imaging. Confocal images were acquired immediately using a Nikon A1R microscope equipped with a 60× oil-immersion objective. For immunostaining of acetylated α-tubulin and detyrosinated tubulin, the same protocol was followed except that cells were exposed to pulsed blue light illumination (470 nm, 40 μW/mm^2^, 5 sec on and 30 sec off) for 0, 10, 30, or 60 min for light-induced α-tubulin acetylation or 0, 20, 60, or 90 min for light-induced tubulin detyrosination prior to fixation.

### Confocal imaging and image analysis

Fluorescence imaging was mainly performed on a Nikon Eclipse Ti-E microscope equipped with an A1R-A1 confocal module with LU-N4 laser sources (argon-ion: 405 and 488 nm; diode: 561 nm), CFI (chrome-free infinity) plan Apochromat VC series objective lenses (60 × oil or 40 × oil), and a live cell culture cage to main the temperature at 37 °C with 5% CO_2_. Photoactivation with persistent illumination or pulses of repeated dark-light cycles was achieved by an external blue light source (470 nm, 40 µW/mm^2^, ThorLabs Inc., Newton, NJ, USA) or by utilizing the built-in 488-nm excitation channel (5% input or at a power density of 10 µW/mm^2^). In some experiments, images were acquired using a a DeltaVision imaging workstation (GE Healthcare) equipped with a 100 ×/1.45 oil lens and a CoolSNAP EMCCD camera to achieve higher temporal resolution.

All the acquired images were analyzed using the NIS-Elements imaging software (version 4.5 1.00), and the results were plotted using the Prism 10.5.0 (673) software (GraphPad). Cytosolic fluorescence intensities were quantified using the semi- or fully-automatic image analysis tool in the NIS-Elements software package. Regions of interest (ROIs), such as MT, MT plus-ends (comets), and adjacent cytosolic areas, were defined and measured using the “Intensity Line Profile” tool by drawing a line to extract fluorescence distribution profiles. The fluorescence intensity of the target protein within each ROI was quantified for further statistical analysis. Typically, 6–8 regions per cell were analyzed to obtain averaged values

### Cell cycle and viability analysis

24 h post-transfection, HeLa cells expressing either mCh-OptoMT or mCh-CRY2 (as control) were washed with PBS and fixed in ice-cold 70% ethanol at 4°C for 30 min. After washing with PBS, the cells were incubated with 1 mg/ml of DAPI (Sigma D9542) and analyzed by flow cytometry. Histograms of cell cycle distribution were acquired using a BD LSRII flow cytometer (BD Biosciences), and the proportions of cells in G_0_/G_1_, S, and G_2_/M phases were determined using FlowJo v10 software. Each sample was assayed in triplicate. Cell viability was assessed by the standard trypan blue staining assay as described previously [98].Cell viability was assessed by the standard trypan blue staining assay as described previously [98].

### Western blot analysis

On day 1, cells were trypsinized and washed three times with ice-cold PBS, then lysed directly in RIPA buffer supplemented with 1× protease inhibitor cocktail and phosphatase inhibitor cocktail for 30 min on ice. Lysates were centrifuged, and the resulting supernatant was transferred to new tubes and denatured at 95 °C for 5 min in 1× SDS loading buffer (100 mM Tris-HCl, 4% SDS, 0.2% bromophenol blue, 20% glycerol, 200 mM DTT, pH 7.4). Equal amounts of protein were resolved on a 10% SDS-PAGE gel and transferred onto nitrocellulose membranes. Membranes were incubated with the indicated primary antibodies overnight at 4 °C. On day 2, the membranes were washed and incubated with secondary antibodies for 1 h at RT and visualized using the ChemiDoc Imaging System (Bio-Rad) with West-Q Pico Dura ECL substrate (GenDEPOT). Densitometric analysis of immunoblot bands was performed using the Gel Analysis function in Image J (NIH; version 2.16.0/1.54p). Band intensities were quantified after background subtraction and normalized to GAPDH or the indicated reference proteins.

### Real-time intracellular Ca^2+^ measurements

Ca^2+^ influx was monitored in HeLa cells stably expressing the green Ca^2+^ indicator GCaMP6s and transiently transfected with either mCherry-OptoTIP or mCh-OptoTIP-SAW. Time-lapse fluorescence imaging was performed with 10 s intervals, and images were analyzed using NIS-Elements AR software (version 4.5 1.00) with manually defined regions of interest. To measure store-operated Ca^2+^ entry (SOCE), cells were incubated in Ca^2+^ free HBSS buffer (107 mM NaCl, 7.2 mM KCl, 1.2 mM MgCl_2_, 11.5 mM glucose and 20 mM HEPES-NaOH, pH 7.2) 5 min prior to imaging. ER Ca^2+^ stores were depleted by using 1 μM thapsigargin (TG), followed by addition of CaCl_2_ to a final concentration of 2 mM to induce Ca^2+^ influx.

### *C. elegans* strains and handling

All strains were derived from the N2 Bristol wild type (see strain information in **Supplementary Table 2**). Animals were cultured at 20°C on 5 cm NG agar plates with OP50 *E. coli* as food [99]. Standard *C. elegans* nomenclature was followed [100]. To provide a non-fluorescent selectable marker for transgenesis, the mutation *dpy-20(e1362)* was crossed into the strain OD2765, which harbors the *itSi916* single-copy transgenic insertion expressing GFP::TBB-2 [101]. Rescuing *dpy-20(*+) plasmid pMH86 was co-injected with the respective plasmid encoding mCherry-OptoMT (*dpy-7p>mCherry::OptoMT*) or mCherry-OptoTIP (*dpy-7p>mCherry::OptoTIP*) to generate transgenic extrachromosomal arrays; non-Dpy progreny co-expressed mCherry::OptoMT or mCherry::OptoTIP in epithelial cells expressing GFP::TBB-2 (β-tubulin).

### Statistical analysis

All data are presented as mean ± sem unless otherwise noted. Sample sizes (n) were listed for each experiment. For Opto-αTAT1 and SxIP tag-related experiments, two-tailed Student’s t-test was used to analyze significant differences between different groups. For blue light-induced dimerization and oligomerization, two-tailed Student’s t-test was used to analyze significant differences. For all statistics, ns, P ≥ 0.05; ***P < 0.001.

### Data availability

The datasets generated and/or analyzed during this study are available as Supplementary Information. Source data for are also available upon request.

## ACKNOWLEDGEMETNS

This work was supported by the the National Institutes of Health (R01GM144986, and R21AI174606 to Y.Z.; R35GM144237 to DJR; R01CA240258, R01DK132286, and R35HL166557 to Y.H.). We thank Shaohe Wang and Karen Oegema for sharing the unpublished C. elegans strain OD2765. Some strains were provided by the Caenorhabditis Genetics Center, which is funded by the NIH Office of Research Infrastructure Programs (P40 OD010440).

## AUTHOR CONTRIBUTIONS

YZ, YH and GM conceived the ideas and directed the work. GM and YZ designed the study. GM, XL, MC, DD, TL and YZ designed and generated all the plasmid constructs. GM, XL, YH and YZ developed and characterized the optogeneic tools. GM, TD and DR performed *C. elgans* studies. GM, YZ, XL and TL extended the applications of tools. GM, XL, TL and YZ analyzed data. GM, XL, TL, and YZ wrote the manuscript, with inputs from all other co-authors.

## COMETING FINANCIAL INTERESTS

The authors declare no competing financial interests.

