## Supplemental figures for "A single-component optogenetic toolkit for programmable control of microtubule"

**The PDF file contains:**

Supplementary Figures 1-9

Supplementary Table 2

Supplementary Movies 1-15

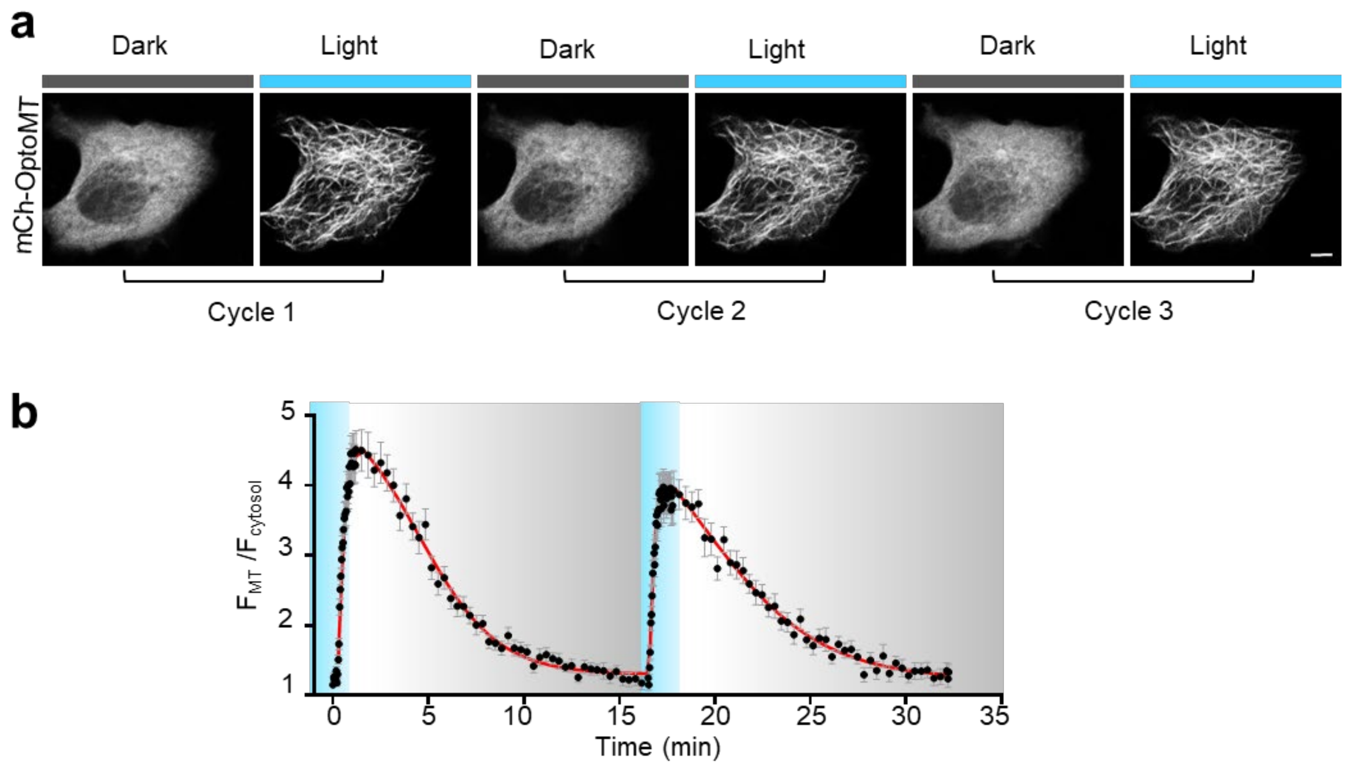


**Supplementary Figure 1 | OptoMT enables rapid and reversible labeling of MTs.** (Related to Fig. 1).

(a) Confocal images of HeLa cells expressing mCh-OptoMT during three repeated dark-light cycles (470 nm, 40 µW/mm$^2$), showing reversible labeling of MTs. Images for the first two cycles were also shown in **Figure 1e**. Scale bar, 5 µM.

(b) Quantitative analysis of MT-to-cytosol fluorescence intensity ratio of mCh-OptoMT in HeLa cells, showing OptoMT-mediated reversible labeling of MTs during two repeated light-dark cycles (470 nm, 40 µW/mm$^2$). Data were fit by a single exponential decay function ($t_{1/2, on}$ = 10.1 ± 4.2 sec; $t_{1/2, off}$ = 210 ± 28 sec). Data are presented as mean ± sem. n = 37 cells from three independent biological replicates. Also see **Supplementary Video 2**.

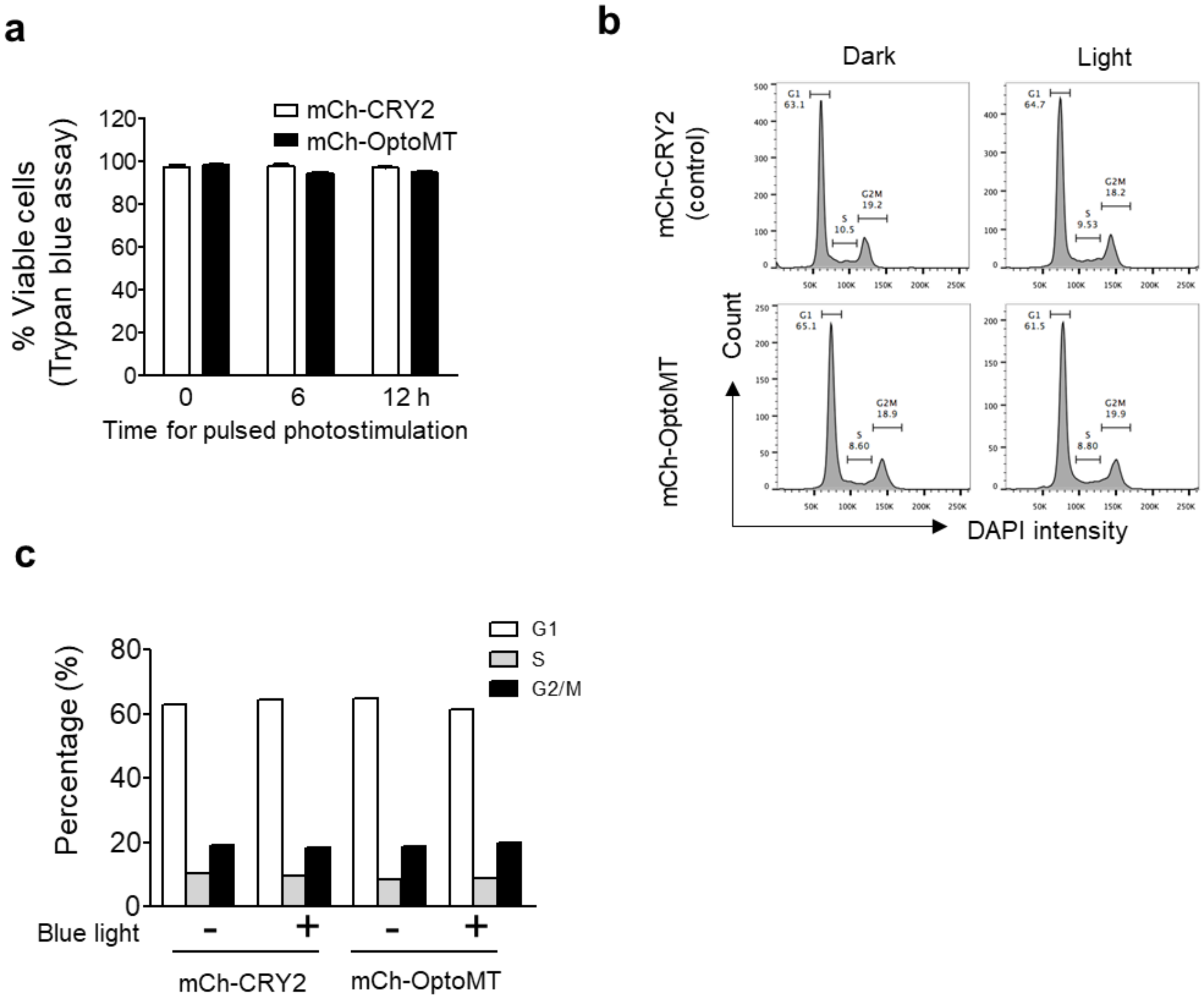


**Supplementary Figure 2 | mCh-OptoMT expression in HeLa cells does not significantly affect cell cycle progression, cell division, and cell viability.** (Related to Fig. 1).

(a) Quantification of cell viability. HeLa cells expressing either mCh-CRY2-PHR (control) or mCh-OptoMT were maintained in the dark (0 h) or exposed to pulsed blue light (3 s per min for 6 h or 12 h). Data are presented as mean ± sem. n = 3 independent biological replicates.

(b) Cell cycle analysis of HeLa cells expressing either mCh-OptoMT or mCh-CRY2 (control), with or without pulsed blue light exposure (3 s per min, 470 nm, 40 µW/mm$^2$).

(c) Quantification of cell populations at each representative cell cycle stage, determined by flow cytometry analysis of HeLa cells expressing mCh-OptoMT or mCh-CRY2 (control).

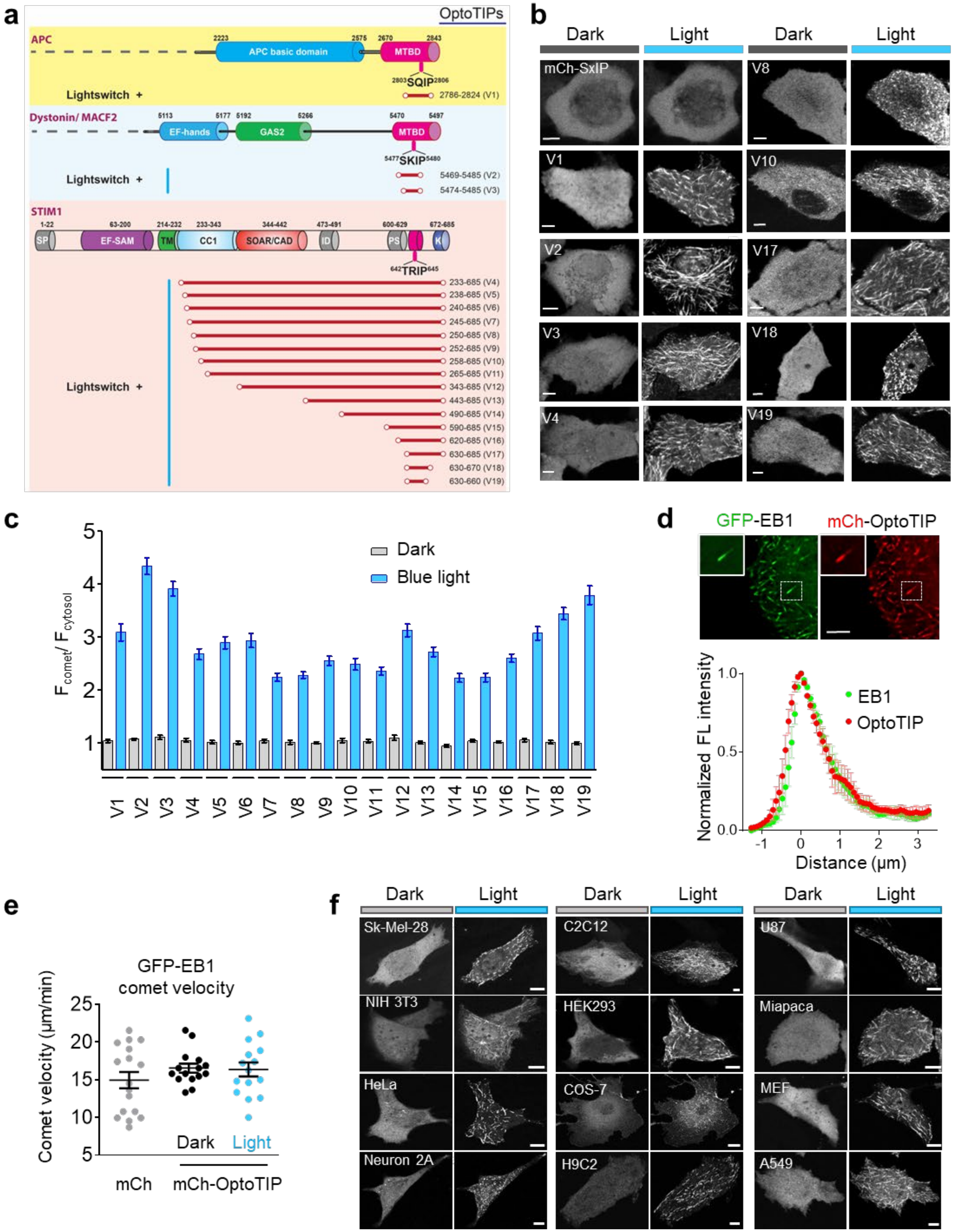
a
OptoTIPs
APC
APC basic domain
MTBD
SQIP
Lightswitch +
2786-2824 (V1)
Dystonin/ MACF2
EF-hands
GAS2
MTBD
SKIP
Lightswitch +
5469-5485 (V2)
5474-5485 (V3)
STIM1
SP
EF-SAM
TM
CC1
SOAR/CAD
ID
PS
K
TRIP
Lightswitch +
233-685 (V4)
238-685 (V5)
240-685 (V6)
245-685 (V7)
250-685 (V8)
252-685 (V9)
258-685 (V10)
265-685 (V11)
343-685 (V12)
443-685 (V13)
490-685 (V14)
590-685 (V15)
620-685 (V16)
630-685 (V17)
630-670 (V18)
630-660 (V19)
b
Dark
Light
Dark
Light
mCh-SxIP
V1
V2
V3
V4
V8
V10
V17
V18
V19
c
Dark
Blue light
$F_{comet}/F_{cytosol}$
V1 V2 V3 V4 V5 V6 V7 V8 V9 V10 V11 V12 V13 V14 V15 V16 V17 V18 V19
d
GFP-EB1
mCh-OptoTIP
Normalized FL intensity
EB1
OptoTIP
Distance (μm)
e
GFP-EB1
comet velocity
Comet velocity (μm/min)
Dark
Light
mCh
mCh-OptoTIP
f
Dark
Light
Dark
Light
Dark
Light
Sk-Mel-28
NIH 3T3
HeLa
Neuron 2A
C2C12
HEK293
COS-7
H9C2
U87
Miapaca
MEF
A549

**Supplementary Figure 3 | Characterization of engineered mCh-OptoTIP constructs.** (Related to Fig. 2).

(a) EB1-binding SxIP motifs derived from APC, dystonin (DST or MACF2) and STIM1 were fused to the photosensory domain of CRY2 to generate 19 OptoTIP variants (V1 to V19). Abbreviations: APC, adenomatous polyposis coli; MTBD, microtubule-binding domain; DST, dystonin; GAS2, growth arrest-specific 2; STIM1, stromal interaction molecule 1; SP, signal peptide; EF-SAM, EF-hand and sterile alpha motif; TM, transmembrane domain; CC1, coiled-coil; SOAR or CAD, STIM1 Orai-activating region or CRAC-activating domain; ID, inhibitory domain; PS, proline-serine rich sequence; K, polybasic lysine-rich domain.

(b) Confocal images of HeLa cells expressing the indicated mCh-OptoTIP variants, acquired before (dark bar) and after 30 s blue light stimulation (blue bar). Scale bar, 5 µM.

(c) Quantification of the comet-to-cytosol fluorescence intensity ratio of mCh-OptoTIP variants in HeLa cells shown in panel (b), measured before and after 30 s blue light stimulation. Data are presented as mean ± sem from at least 20 cells from three independent biological replicates.

(d) Top, confocal images of HeLa cells co-expressing GFP-EB1 and mCh-OptoTIP under blue light illumination. Bottom, fluorescence intensity line profiles showing tight colocalization of EB1-GFP and mCh-OptoTIP comets. Scale bar, 5 µM. Data are presented as mean ± sem. n = 58 comets from three independent biological replicates.

(e) Quantification of comet velocity in HeLa cells expressing either mCh (control) or mCh-OptoTIP with or without photostimulation. Blue light-dependent labeling of mCh-OptoTIP at MT plus-ends did not significantly alter the comet velocity of GFP-EB1. Data are presented as mean ± sem from at least 15 cells from three independent biological replicates.

(f) Confocal images of mCh-OptoTIP (V2) expressed in multiple cell lines. All tested cell types exhibited blue light-inducible MT comet labeling without the need of EB1 overexpression. Scale bar, 5 µM.

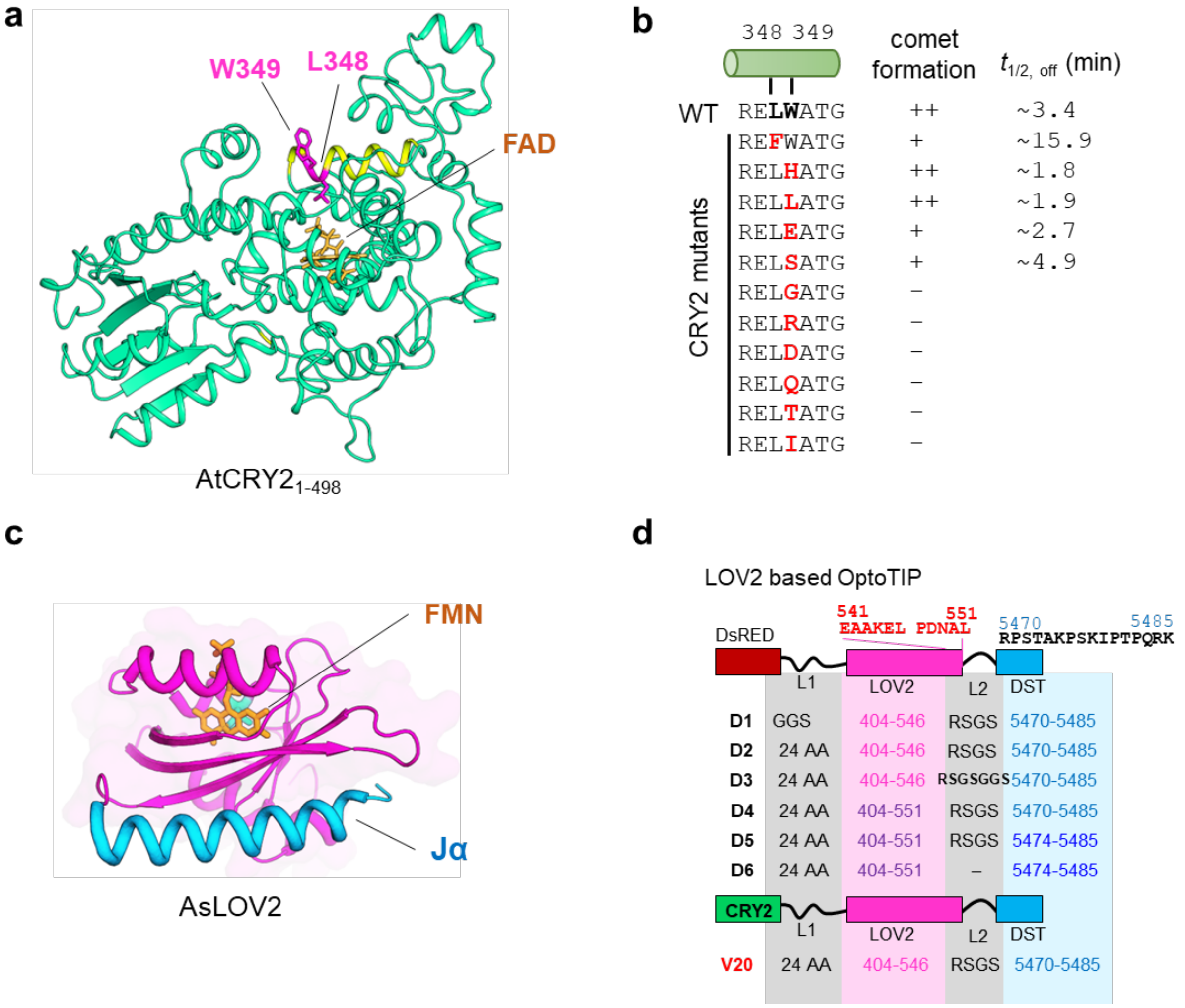


**Supplementary Figure 4 | Tuning the kinetics of mCh-OptoTIP variants**. (Related to Fig. 2).

(a) The 3D modeled structure of the N-terminal photolyase homology region (PHR) domain of *Arabidopsis* CRY2. Mutations at residues L348 and W349 (magenta sticks) are approximately 10 Å from the flavin adenine dinucleotide (FAD, yellow stick) cofactor binding site.

(b) CRY2 was engineered with fast (L348F) or slow photocycle (W349X) mutations to modulate off-kinetics of light-induced activation.

(c) The crystal structure of the dark-state AsLOV2 protein (PDB: 2V0W), highlighting the core PAS domain (red), C-terminal Jα helix (blue), and the flavin mononucleotide cofactor (FMN, brown).

(d) Domain architecture of LOV2-SxIP fused either to tetrameric DsRed (variants D1-D6) or the light-inducible oligomerization module CRY2 (V20). The EB1-binding SxIP motif (derived from DST, residues 5469-5485) was appended to the C-terminus of the LOV2 domain and subsequently fused downstream of DsRed or CRY2, enabling light-dependent modulation of MT plus-end tracking.

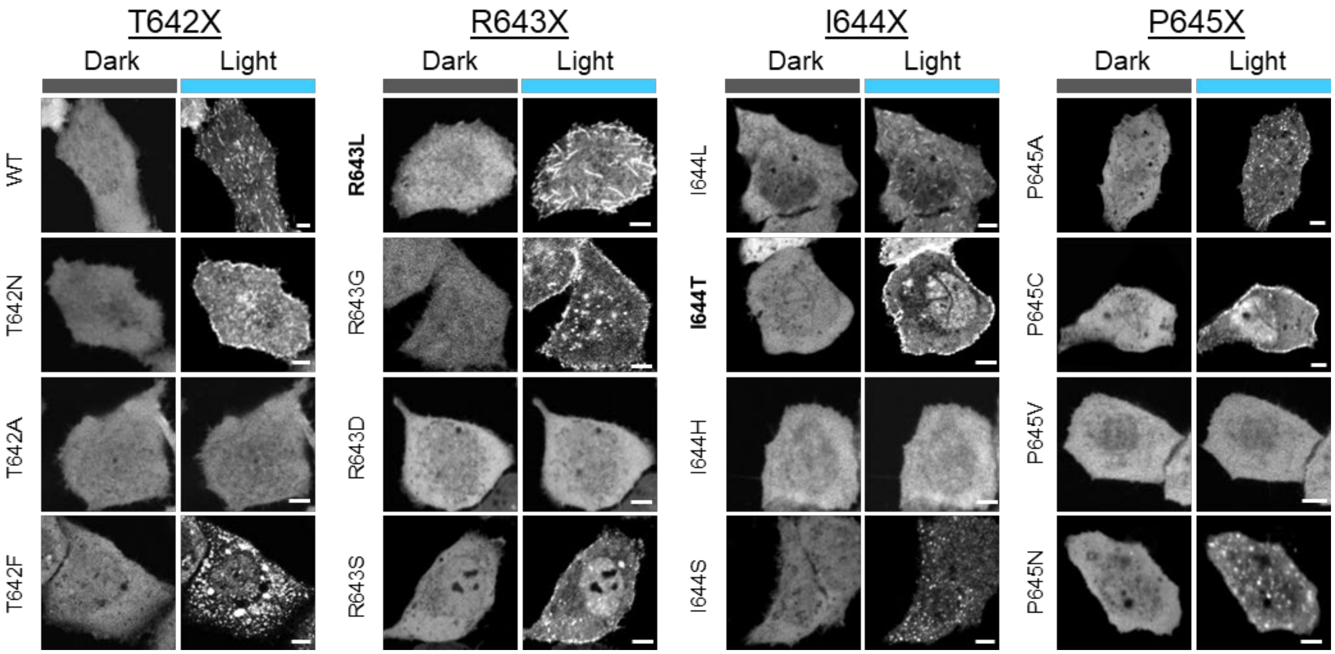


**Supplementary Figure 5 | Live-cell imaging of OptoTIP-V17 variants before and after photostimulation.** (Related to Fig. 3).

Confocal images of HeLa cells expressing mCh-OptoTIP-V17 (CRY2-STIM1 630-685) variants, in which the EB1-binding TRIP motif (residues 642-645) was individually substituted with each of 19 other amino acids. Based on their localization patterns before and after bule light illumination, the mutants exhibited four distinct phenotypes: (i) MT plus-end tracking via EB1 binding, (ii) PM translocation mediated by interactions with phosphoinositides, (iii) uniform cytosolic distribution, or (iv) punctate clustering resulting from CRY2 homo-oligomerization. Scale bar, 5 µm.

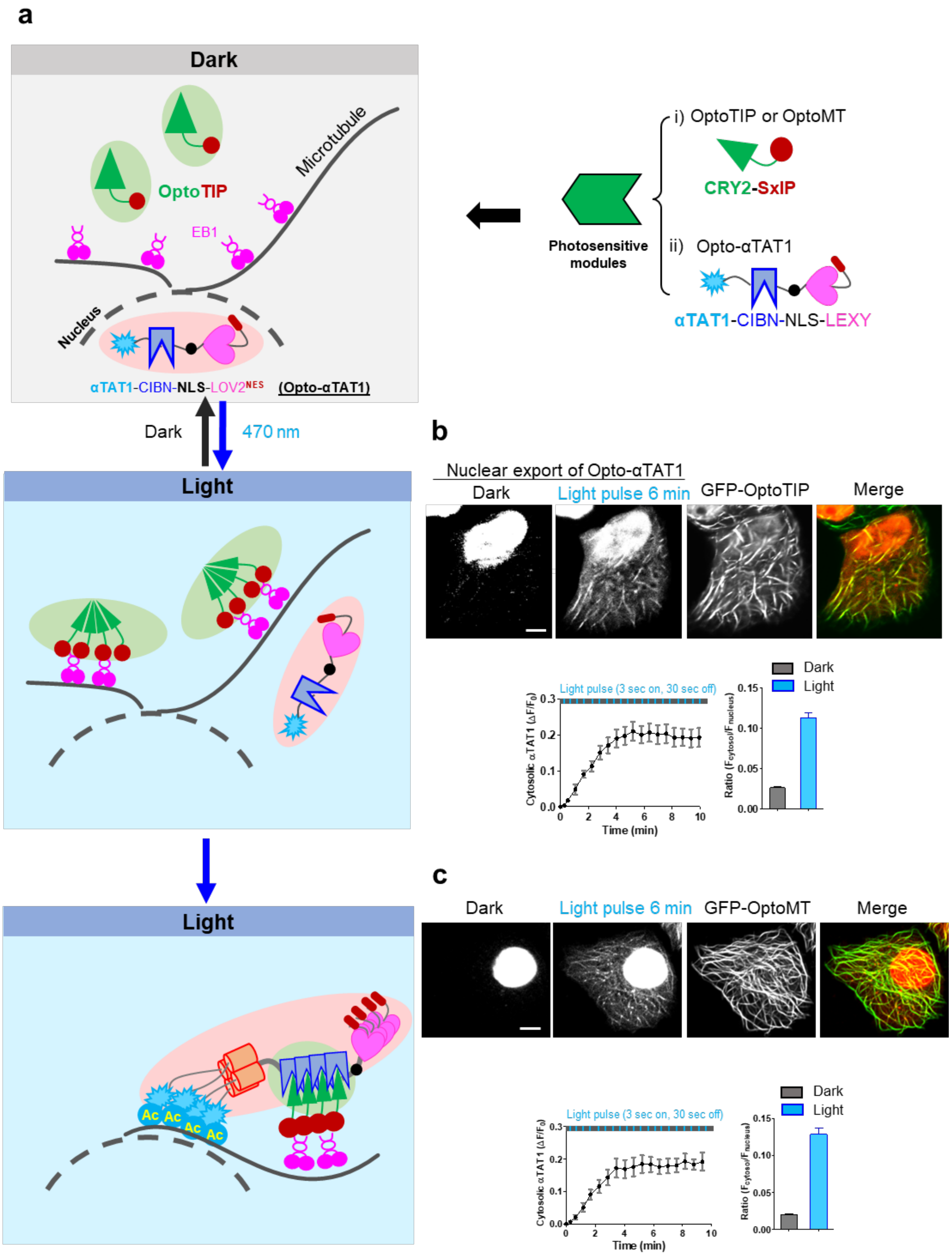
a
Dark
OptoTIP
Microtubule
EB1
Nucleus
αTAT1-CIBN-NLS-LOV2NES (Opto-αTAT1)
Photosensitive modules
i) OptoTIP or OptoMT
CRY2-SxIP
ii) Opto-αTAT1
αTAT1-CIBN-NLS-LEXY
Dark
470 nm
Light
Light
Ac
b
Nuclear export of Opto-αTAT1
Dark
Light pulse 6 min
GFP-OptoTIP
Merge
Light pulse (3 sec on, 30 sec off)
Cytosolic αTAT1 (ΔF/F0)
Time (min)
Dark
Light
Ratio (Fcytosol/Fnucleus)
c
Dark
Light pulse 6 min
GFP-OptoMT
Merge
Light pulse (3 sec on, 30 sec off)
Cytosolic αTAT1 (ΔF/F0)
Time (min)
Dark
Light
Ratio (Fcytosol/Fnucleus)

**Supplementary Figure 6 | Design of Opto-αTAT1 and its integration with OptoTIP or OptoMT to enable light-inducible tubulin acetylation.** (Related to Fig. 4).

(a) Schematic illustrating the design of Opto-αTAT1 and its integration with OptoTIP or OptoMT.

Top right: a simplified diagram of Opto-αTAT1, which integrates two photosensitive components for precise control of αTAT1 nuclear export and MT-targeting. The LEXY nuclear export system enables blue light-induced translocation of αTAT1 from the nucleus to the cytosol (αTAT1-mCh-CIBN-NLS-LEXY). Once exported to the cytosol, αTAT1 is further recruited to MTs via light-dependent heterodimerization with CRY2-containing OptoTIP or OptoMT.

Left: schematic showing the sequential mechanism by which Opto-αTAT1 and OptoTIP coordinate to induce MT acetylation. In the dark, Opto-αTAT1 remains sequestered in the nucleus due to LOV2-mediated caging of the nuclear export signal (NES. Upon blue light illumination, NES exposure triggers cytosolic translocation of Opto-αTAT1. Simultaneously, photostimulation induces OptoTIP or OptoMT binding to MT filaments or MT plus-ends, respectively, which in turn recruits Opto-αTAT1 via CRY2-CIBN heterodimerization, thereby positioning the enzyme for targeted tubulin acetylation.

(b-c) Confocal images showing light-induced nuclear export of mCh-Opto-αTAT1 (red) and its subsequent co-localization with GFP-OptoTIP (b, green) or GFP-OptoMT (c, green). The corresponding time courses and quantification of nuclear export were shown on the bottom. Export kinetics were monitored for up to 10 minutes, with cytosolic mCh intensity peaking around 5-6 min. Although a substantial portion of Opto-αTAT1 remained in the nucleus, the cytosolic fraction was sufficient to induce effective tubulin acetylation (see **Fig. 4e-j**). Scale bar, 5 µm. Data were presented as mean ± sem. n= 8 cells from three independent biological replicates for the time-course curves; n= 42 (OptoTIP) and 29 (OptoMT) cells for fluorescence intensity quantification from three independent biological replicates.

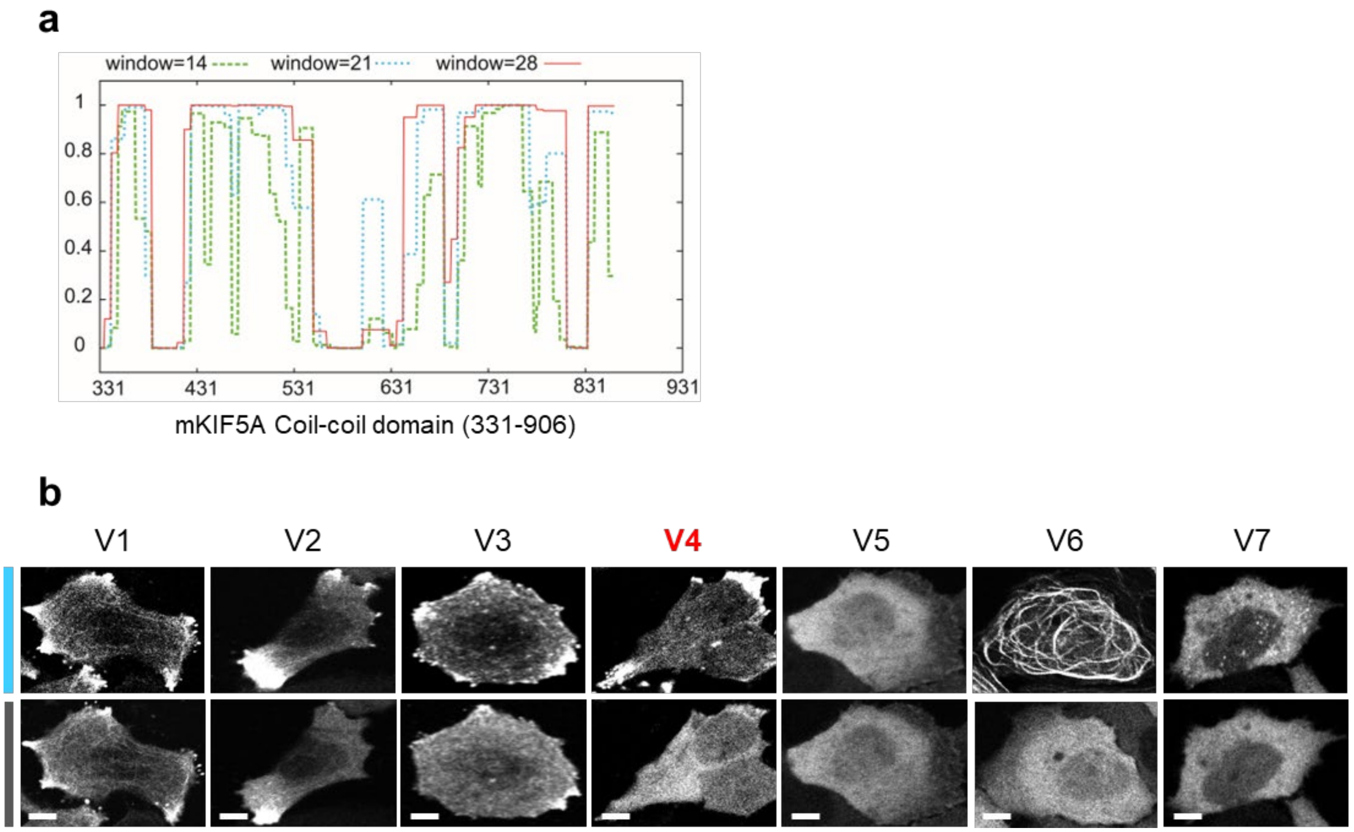


**Supplementary Figure 7 | Design and screening of mKIF5A-based OptoMotor variants**. (Related to Fig. 5).

(a) Predicted coiled-coil probability along the KIF5A stalk region (residues 331-906) calculated using sliding windows of 14, 21, and 28 in the COILS algorithm.

(b) Confocal images of HeLa cells expressing the indicated OptoMotor variants, acquired before and after blue light illumination (470 nm, 40 µW/mm²). V1-V3 exhibited constitutive peripheral accumulation, V4 showed robust light-induced peripheral redistribution, V5 failed to respond, V6 labeled MTs, and V7 formed puncta without MT engagement. Scale bar, 5 µm.

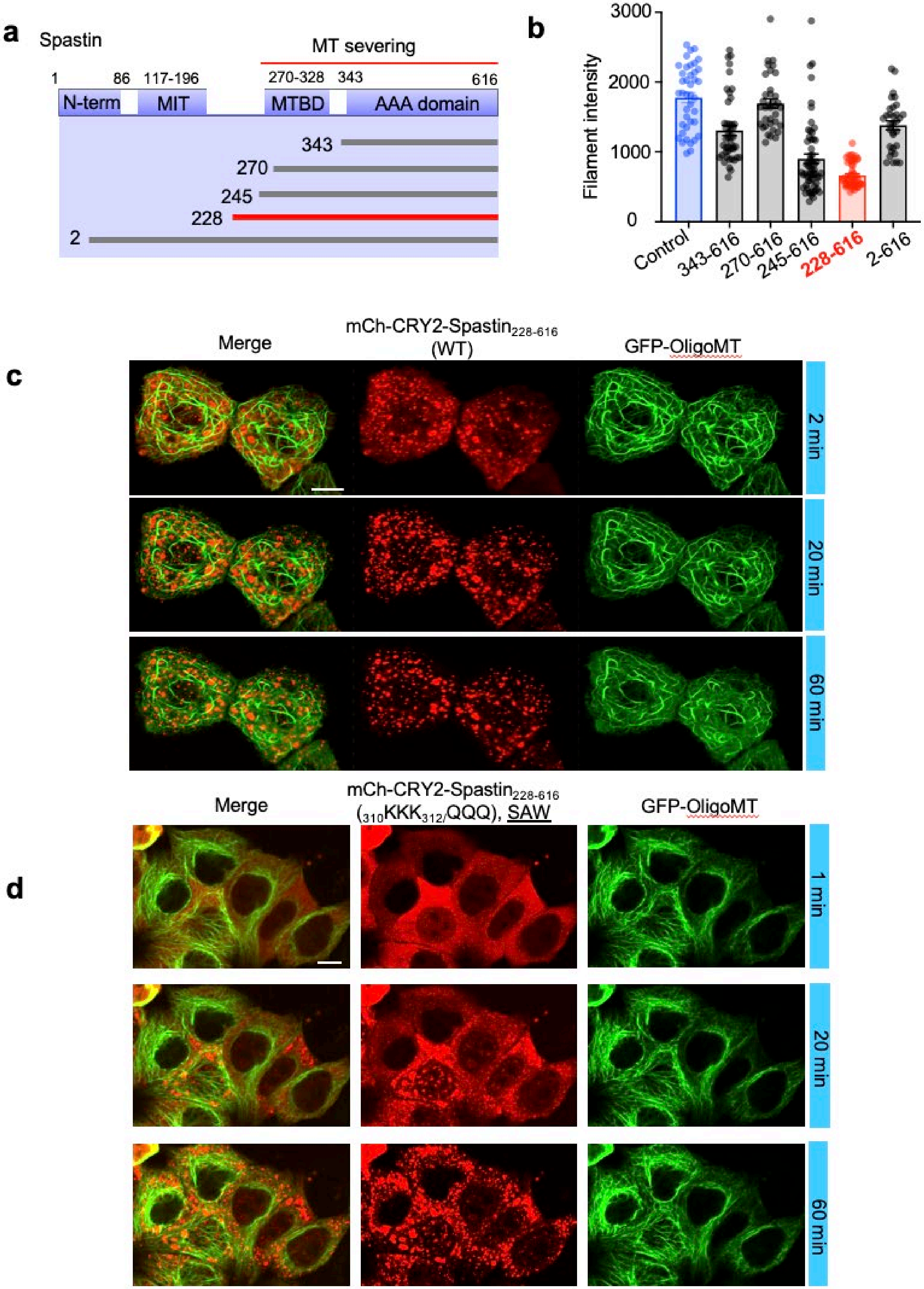
a
Spastin
MT severing
1
86
117-196
270-328
343
616
N-term
MIT
MTBD
AAA domain
343
270
245
228
2
b
3000
2000
1000
0
Filament intensity
Control
343-616
270-616
245-616
228-616
2-616
Merge
mCh-CRY2-Spastin228-616 (WT)
GFP-OligoMT
c
2 min
20 min
60 min
Merge
mCh-CRY2-Spastin228-616 (310KKK312/QQQ), SAW
GFP-OligoMT
d
1 min
20 min
60 min

**Supplementary Figure 8 | Mapping the MT-severing module in spastin.** (Related to Fig. 6).

(a) Domain architecture of the MT-severing protein spastin. Each truncation fragment was fused to the C-terminus of mCh-OptoMT to evaluate light-inducible MT severing activity.

(b) Quantification of MT filament integrity (indicated by GFP-OligoMT fluorescence intensity) in HeLa cells co-expressing GFP-OligoMT and the indicated variants 6 h of blue light exposure (470 nm, 40 μW/mm²) using 3 s on and 30 s off pulses. Data are presented as mean ± sem from at least 10 cells from three independent biological replicates.

(c-d) Comparison of basal severing activity between WT $spastin_{228\text{-}616}$ and the mutant, K3/Q3 (SAW). Shown are time-lapse confocal imaging of HeLa cells co-expressing GFP-OligoMT (green) and either (c) mCherry-CRY2-$spastin_{228\text{-}616}$ (red) or (d) mCh-CRY2- $spastin_{228\text{-}616}$ K3/Q3 mutant (SAW) (red) during 60 min of blue light illumination. Scale bar, 5 μm.

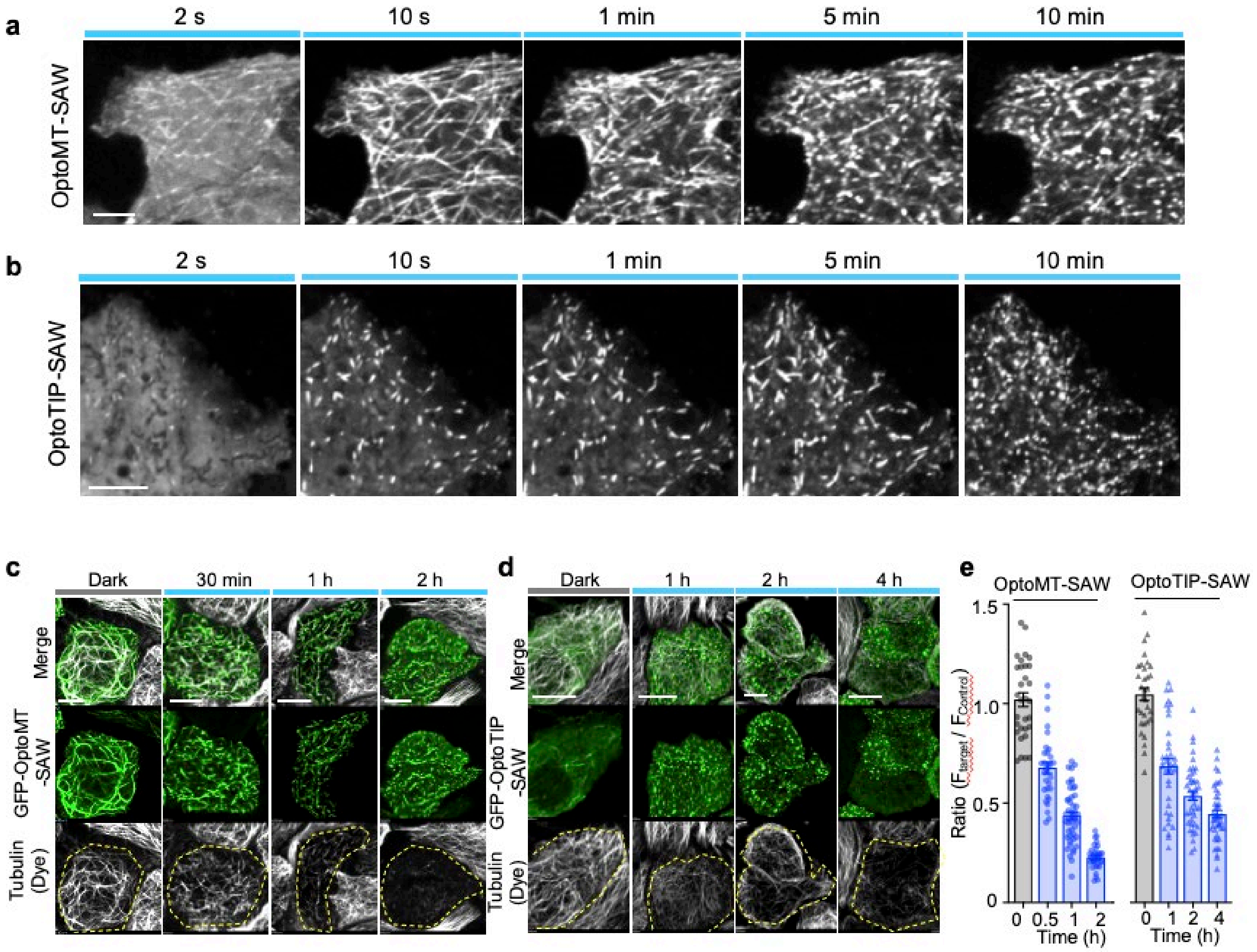


**Supplementary Figure 9 | Real-time imaging of OptoMT-SAW and OptoTIP-SAW**. (Related to Fig. 7).

(a) Time-lapse confocal imaging of HeLa cells expressing mCh-OptoMT-SAW under blue light stimulation. Upon illumination, mCh-OptoMT-SAW rapidly decorated MTs and subsequently mediated filament severing, leading to the fragmentation of long MTs into shorter segments and the formation of punctate clusters after 10 min blue light illumination. Scale bar, 5 µm.

(b) Time-lapse confocal imaging of HeLa cells expressing mCh-OptoTIP-SAW under blue light stimulation. Upon illumination, mCh-OptoTIP-SAW localized to growing MT plus-ends, forming characteristic comet-like structures. Continued light exposure induced MT severing events, with puncta clusters emerging after approximately 10 min. Scale bar, 5 µm.

(c-e) Confocal images of HeLa cells expressing (c) GFP-OptoMT-SAW or (d) GFP-OptoTIP-SAW. Cells were exposed to blue light illumination (470 nm, 40 µW/mm²) for varying durations and subsequently stained with a deep-red tubulin tracker. The quantification of the data as a function of photostimulation time is shown on the bar graph. Scale bar, 5 µm. Data are presented as mean ± sem. n = 30-40 cells from three independent biological replicates.

**Supplementary Table 1 |** Summary of activation and deactivation half-lives of tools in the study.

| | Version | Components | $F_{MT}/F_{cytosol}$ or $F_{comet}/F_{cytosol}$ | | On phase | Off phase |
|---|---|---|---|---|---|---|
| | | | Dark | Light | ($t_{1/2}$, sec) | ($t_{1/2}$, sec) |
| **OptoMT** | V1 | CRY2$_{1\text{-}498}$-KIF5A$_{1\text{-}330}$ | 1.1 | 3.5 | | |
| | **V2** | CRY2$_{1\text{-}498}$-EB1$_{1\text{-}191}$ | 1.1 | 3.5 | | |
| | V3 | **CRY2$_{1\text{-}498}$-Clip170$_{129\text{-}350}$** | **1.0** | **4.7** | **10.1 ± 4.2** | **210.0 ± 28.2** |
| | V4 | CRY2$_{1\text{-}498}$-CAMSAP1$_{1270\text{-}1473}$ | 1.7 | 4.9 | | |
| **OptoTIP variants** | V1 | CRY2$_{1\text{-}498}$-APC$_{2786\text{-}2824}$ | 1.0 | 3.0 | | |
| | **V2** | **CRY2$_{1\text{-}498}$-DST$_{5469\text{-}5485}$** | **1.0** | **4.4** | **11.1 ± 4.7** | **204.0 ± 43.8** |
| | | CRY2$_{1\text{-}498,\ L348F}$-DST$_{5469\text{-}5485}$ | 1.0 | 3.2 | | ~ 954 |
| | | CRY2$_{1\text{-}498,\ W349H}$-DST$_{5469\text{-}5485}$ | 1.0 | 4.2 | | ~ 108 |
| | | CRY2$_{1\text{-}498,\ W349L}$-DST$_{5469\text{-}5485}$ | 1.0 | 4.5 | | ~ 114 |
| | | CRY2$_{1\text{-}498,\ W349E}$-DST$_{5469\text{-}5485}$ | 1.0 | 3.5 | | ~ 162 |
| | | CRY2$_{1\text{-}498,\ W349A}$-DST$_{5469\text{-}5485}$ | 1.0 | 1.1 | | |
| | | CRY2$_{1\text{-}498,\ W349G}$-DST$_{5469\text{-}5485}$ | 1.0 | 1.2 | | |
| | | CRY2$_{1\text{-}498,\ W349S}$-DST$_{5469\text{-}5485}$ | 1.0 | 1.1 | | |
| | | CRY2$_{1\text{-}498,\ W349M}$-DST$_{5469\text{-}5485}$ | 1.0 | 1.5 | | |
| | | CRY2$_{1\text{-}498,\ W349R}$-DST$_{5469\text{-}5485}$ | 1.0 | 1.2 | | |
| | | CRY2$_{1\text{-}498,\ W349D}$-DST$_{5469\text{-}5485}$ | 1.0 | 1.1 | | |
| | | CRY2$_{1\text{-}498,\ W349Q}$-DST$_{5469\text{-}5485}$ | 1.0 | 1.4 | | |
| | | CRY2$_{1\text{-}498,\ W349T}$-DST$_{5469\text{-}5485}$ | 1.0 | 1.1 | | |
| | | CRY2$_{1\text{-}498,\ W349I}$-DST$_{5469\text{-}5485}$ | 1.0 | 1.3 | | |
| | V3 | CRY2$_{1\text{-}498}$-DST$_{5474\text{-}5485}$ | 1.0 | 4.3 | | |
| | V4 | CRY2$_{1\text{-}498}$-STIM1$_{233\text{-}685}$ | 1.0 | 2.6 | ~ 45 | ~ 320 |
| | V5 | CRY2$_{1\text{-}498}$-STIM1$_{238\text{-}685}$ | 1.0 | 2.9 | | |
| | V6 | CRY2$_{1\text{-}498}$-STIM1$_{240\text{-}685}$ | 1.0 | 2.9 | | |
| | V7 | CRY2$_{1\text{-}498}$-STIM1$_{245\text{-}685}$ | 1.0 | 2.2 | | |
| | V8 | CRY2$_{1\text{-}498}$-STIM1$_{250\text{-}685}$ | 1.0 | 2.3 | | |
| | V9 | CRY2$_{1\text{-}498}$-STIM1$_{252\text{-}685}$ | 1.0 | 2.5 | | |
| | V10 | CRY2$_{1\text{-}498}$-STIM1$_{258\text{-}685}$ | 1.0 | 2.4 | | |
| | V11 | CRY2$_{1\text{-}498}$-STIM1$_{265\text{-}685}$ | 1.0 | 2.3 | | |
| | V12 | CRY2$_{1\text{-}498}$-STIM1$_{343\text{-}685}$ | 1.0 | 3.1 | | |
| | V13 | CRY2$_{1\text{-}498}$-STIM1$_{443\text{-}685}$ | 1.0 | 2.7 | | |
| | V14 | CRY2$_{1\text{-}498}$-STIM1$_{490\text{-}685}$ | 1.0 | 2.2 | | |
| | V15 | CRY2$_{1\text{-}498}$-STIM1$_{590\text{-}685}$ | 1.0 | 2.2 | | |
| | V16 | CRY2$_{1\text{-}498}$-STIM1$_{620\text{-}685}$ | 1.0 | 2.6 | | |
| | V17 | CRY2$_{1\text{-}498}$-STIM1$_{630\text{-}685}$ | 1.0 | 3.0 | ~ 15 | ~ 260 |
| | V18 | CRY2$_{1\text{-}498}$-STIM1$_{630\text{-}670}$ | 1.0 | 3.4 | | |
| | V19 | CRY2$_{1\text{-}498}$-STIM1$_{630\text{-}660}$ | 1.0 | 3.8 | ~ 12 | ~ 220 |
| | V20 | **CRY2$_{1\text{-}498}$-LOV2-DST$_{5470\text{-}5485}$** | **1.0** | **2.1** | **9.1 ± 4.1** | **40.2 ± 6.6** |
| **Opto-αTAT1** | | αTAT1-CIBN-NLS-LEXY | | | **114 ± 15** | **103 ± 8** |

**Supplementary Table 2 |** Summary of C. elegans strains used in the study.

| Strain | Genotype |
|---|---|
| DV2162 | *dpy-20(e1362*cs*)* IV 5 x outcrossed |
| DV3482 | *itSi916[pOD2050/pSW384; dpy-7p>GFP::tbb-2 3'UTR+Cbr-unc-119(+)]* I; *dpy-20(e1362*cs*)* IV |
| DV3515 | *itSi916[pOD2050/pSW384; dpy-7p>GFP::tbb-2 3'UTR+Cbr-unc-119(+)]* I; *dpy-20(e1362*cs*)* IV; *reEx210[pTD61[dpy-7p>mCherry::OptoTip]+pMH86(dpy-20(+))]* |
| DV3516 | *itSi916[pOD2050/pSW384; dpy-7p>GFP::tbb-2 3'UTR+Cbr-unc-119(+)]* I; *dpy-20(e1362*cs*)* IV; *reEx211[pTD61[dpy-7p>mCherry::OptoMT]+pMH86(dpy-20(+))]* |
| OD2765 | *itSi916[pOD2050/pSW384; dpy-7p>GFP::tbb-2; Cbr-unc-119(+)]* I; *unc-119(ed3)* III |

### SUPPLEMENTARY VIDEOS

**Supplementary Video 1 | 3D reconstruction of MT labeling by OptoMT.**

Z-stack 3D reconstruction of a HeLa cell expressing mCh-OptoMT following blue light stimulation. Upon illumination, OptoMT robustly labeled the MT cytoskeleton, revealing the filamentous network throughout the cell volume.

**Supplementary Video 2 | Reversible, light-induced labeling of the MT cytoskeleton by OptoMT.**

Time-lapse confocal imaging of HeLa cells expressing mCh-OptoMT during two repeated dark-light cycles (470 nm, 4 mW/cm$^2$). OptoMT rapidly associated with MTs upon illumination and subsequently redistributed uniformly throughout the cytoplasm after light withdrawal, demonstrating reversible binding to the MT network.

**Supplementary Video 3 | Spatially confined activation of OptoMT in living cells.**

HeLa cells expressing mCh-OptoMT were subjected to spatially restricted blue light illumination, which induced MT labeling exclusively within the illuminated region of interest (ROI). Non-illuminated regions remained unchanged, demonstrating the high spatial precision of light-dependent OptoMT activation.

**Supplementary Video 4 | Light-inducible and reversible MT labeling by OptoMT in *C. elegans*.**

L3-stage *C. elegans* larvae expressing mCh::OptoMT exhibited a diffuse cytoplasmic distribution in the dark and rapid recruitment to MTs upon blue light exposure. OptoMT binding was reversible, with an activation half-life ($t_{1/2}$, ON) of about 12 s and a decay half-life ($t_{1/2}$, OFF) of approximately 252 s following light withdrawal.

**Supplementary Video 5| Reversible, light-controlled tracking of MT plus-ends by OptoTIP.**

In HeLa cells, mCh-OptoTIP exhibited reversible comet-like accumulation at MT plus-ends under repeated dark-light cycles. Kinetic analysis revealed an activation half-life ($t_{1/2}$, ON) of around 11 s and a decay half-life ($t_{1/2}$, OFF) of about 204 s following light withdrawal.

**Supplementary Video 6 | Light-induced colocalization of OptoTIP with EB1 at dynamic MT plus-ends.**

In HeLa cells, mCh-OptoTIP (red) and GFP-EB1 (green) displayed strong colocalization at growing MT plus-ends following blue light stimulation, confirming that OptoTIP faithfully tracks +TIP dynamics in living cells.

**Supplementary Video 7 | OptoTIP dynamically tracks MT bundles in *C. elegans* larvae.**

L3-stage *C. elegans* larvae co-expressing GFP::tubulin (green in merge) and mCh::OptoTIP (red in merge) were imaged following blue light illumination. OptoTIP rapidly localized to and tracked dynamic MT bundles along the dorsal and ventral epidermal surfaces, demonstrating light-inducible plus-end labeling in a multicellular organism.

**Supplementary Video 8 | Reversible MT plus-end labeling by OptoTIP in *C. elegans* larvae.**

L3-stage *C. elegans* larvae expressing mCh::OptoTIP were subjected to repeated dark-light cycles. OptoTIP rapidly associated with comet-like structures at the MT plus-ends upon blue light illumination and disappeared after light withdrawal, demonstrating robust and reversible plus-end tracking in vivo.

**Supplementary Video 9 | Reversible comet localization of LOV2-caged OptoTIP (V20) in HeLa cells.**

CRY2-fused, LOV2-caged SxIP motif (OptoTIP V20) exhibited rapid and reversible transitions between a diffuse cytosolic distribution and comet-like plus-end localization during repeated dark-light cycles in HeLa cells, with an activation half-life of about 9 s and a deactivation half-life of around 40.2 s.

**Supplementary Video 10 | Light-controlled nuclear export and MT recruitment of Opto-αTAT1 in HeLa cells.**

Opto-αTAT1 underwent rapid and reversible nuclear export upon blue light illumination ($t_{1/2}$, ON = 1.9 min; $t_{1/2}$, OFF = 1.7 min) in HeLa cells. Following export, Opto-αTAT1 was promptly recruited to MTs through CIBN-mediated binding to OptoMT, demonstrating precise spatiotemporal control of enzyme localization and activity.

**Supplementary Video 11** | **Reversible light-induced redistribution of OptoMotor in HeLa cells**.

mCh-tagged OptoMotor showed peripheral accumulation upon blue light illumination and cytosolic redistribution after light withdrawal in HeLa cells, demonstrating reversible, light-dependent control of subcellular localization.

**Supplementary Video 12 | Light-induced MT severing by OptoMT-SAW construct visualized with independent MT marker OligoMT.**

Time-lapse confocal imaging of HeLa cells expressing the mCh-OptoMT-SAW (red) construct showed light-induced fragmentation of the MT network, visualized by the independent MT marker GFP-OligoMT (green). Scale bar, 5 µm.

**Supplementary Video 13 | Light-induced severing of MT plus-end structures by OptoTIP-SAW visualized with the +TIP marker EB1-GFP.**

Time-lapse confocal imaging of HeLa cells co-expressing mCh-OptoTIP-SAW (red) showed the progressive loss of comet-like +TIPs labeled by EB1-GFP (green) upon blue light illumination.

**Supplementary Video 14 | Accelerated formation of STIM1-ORAI1 puncta at ER-PM junctions following light-induced MT disruption.**

HeLa cells stably expressing STIM1-mCh (red) and ORAI1-GFP (green) were transiently transfected with miRFP fused OptoMT-SAW to enable blue light-inducible MT disassembly. After 18 h expression, cells were subjected to periodic blue light illumination (6 s on, 60 s off, 470 nm, 40 µW/mm$^2$) for 2 h, resulting in near-complete MT disruption. Store-operated calcium entry (SOCE) was subsequently triggered by 1 µM thapsigargin (TG), and time-lapse confocal imaging was performed to monitor STIM1-ORAI1 puncta formation at ER-PM junctions.

The top panel shows control cells (without miRFP-OptoMT-SAW expression) with intact MTs, and the middle panel shows cells expressing miRFP-OptoMT-SAW following light-induced MT disassembly. The bottom panel presents quantitative analysis of puncta fluorescence intensity from selected regions (Region 1, green curve, control cell; Region 2, red curve, MT-disrupted cell), demonstrating that STIM1-ORAI1 puncta formed more rapidly in MT-disrupted cells, consistent with accelerated STIM1 activation kinetics. Scale bar, 5µm.

**Supplementary Video 15 | Light-induced impairment of lysosomal motility by OptoTIP-SAW.**

Top, Time-lapse confocal imaging of HeLa cells co-expressing LAMP1-GFP (green) and mCh-OptoTIP-SAW (red). Upon blue light illumination, lysosomal motility decreased progressively, leading to near-complete immobilization after approximately 15 min as a result of light-induced MT disruption.

Bottom, Control HeLa cells expressing LAMP1-GFP alone displayed active, bidirectional lysosome transport along intact MTs under identical imaging conditions.
